# Lack of evidence for increased transcriptional noise in aged tissues

**DOI:** 10.1101/2022.05.18.492432

**Authors:** Olga Ibáñez-Solé, Alex M. Ascensión, Marcos J. Araúzo-Bravo, Ander Izeta

**Author notes:** **For correspondence:** (AI); (MJA-B). These authors contributed equally to this work (Olga Ibáñez-Solé, Alex M. Ascensión).

## Abstract

Aging is often associated with a loss of cell type identity that results in an increase in transcriptional noise in aged tissues. If this phenomenon reflects a fundamental property of aging remains an open question. Transcriptional changes at the cellular level are best detected by single-cell RNA sequencing (scRNAseq). However, the diverse computational methods used for the quantification of age-related loss of cellular identity have prevented reaching meaningful conclusions by direct comparison of existing scRNAseq datasets. To address these issues we created *Decibel*, a Python toolkit that implements side-to-side four commonly used methods for the quantification of age-related transcriptional noise in scRNAseq data. Additionally, we developed *Scallop*, a novel computational method for the quantification of membership of single cells to their assigned cell type cluster. Cells with a greater *Scallop* membership score are transcriptionally more stable. Application of these computational tools to seven aging datasets showed large variability between tissues and datasets, suggesting that increased transcriptional noise is not a universal hallmark of aging. To understand the source of apparent loss of cell type identity associated with aging, we analyzed cell type-specific changes in transcriptional noise and the changes in cell type composition of the mammalian lung. No robust pattern of cell type-specific transcriptional noise alteration was found across aging lung datasets. In contrast, age-associated changes in cell type composition of the lung were consistently found, particularly of immune cells. These results suggest that claims of increased transcriptional noise of aged tissues should be reformulated.

## Introduction

Concomitant to the large repertoire of known age-associated changes at the cellular level, an increase in transcriptional variability is generally assumed to characterize aged cells and tissues (***Nikopoulou et al., 2019; Uyar et al., 2020; Mendenhall et al., 2021; Vijg, 2021***). This phenomenon was first described by Vijg and colleagues as an *age-related increase in transcriptional noise* (***Bahar et al., 2006***), which is still the most commonly used term (***Warren et al., 2007; Enge et al., 2017; Angelidis et al., 2019***). *Transcriptional noise* is here defined as the measured level of variation in gene expression among cells supposed to be identical (***Raser and O’Shea, 2005***). Later, similar findings have been reported as an increase in *identity noise* (***Salzer et al., 2018***), *cell-cell heterogeneity* (***Kimmel et al., 2019***), *cell-to-cell variability* (***Martinez-Jimenez et al., 2019; Ximerakis et al., 2019***), or *loss of cellular identity* in aged tissues (***Solé-Boldo et al., 2020; Izgi et al., 2022***). While all these claims have in common the notion of cells expressing their core transcriptional program or *transcriptomic signature* in a loose way, there are important methodological differences between the published reports that deserve further scrutiny.

Early studies were based on the quantification of the variance associated with the expression of a few pre-selected transcripts by real-time PCR, on bulk cell and tissue samples (***Bahar et al., 2006; Warren et al., 2007***). With the advent of single-cell RNA sequencing (scRNAseq) technologies, whole-transcriptome variability on aged tissues was studied at the single-cell level. A pioneering study on human pancreas by Quake and colleagues found an age-related increase in transcriptional noise specific to pancreatic *β* cells (***Enge et al., 2017***). The authors introduced a definition of transcriptional noise that was based on whole-transcriptome variability: the ratio between biological and technical variation, where the latter was inferred from External RNA Controls Consortium (ERCC) spike-in variability. As ERCC spike-in controls are not included in every scRNAseq experiment, they proposed two alternative methods that were based on the notion of “distance to centroid”: the greater the gene-based distance between cells of the same cell type, the greater the transcriptional noise associated with them. One of them measured the Euclidean distance to the cell type mean per individual, using the whole transcriptome. The other one measured the Euclidean distance between each cell and the tissue mean, using a set of invariant genes. Soon after, loss of identity was reported in aged murine dermal fibroblasts by measuring the coefficient of variation of the distances between each highly variable gene between the two main fibroblast clusters (***Salzer et al., 2018***). Similar findings were published in early activated CD4+ T cells, based on the observation that the fraction of cells that expressed the core activation program was lower in old animals (***Martinez-Jimenez et al., 2019***). A study on murine aging lung found an increase in cellular heterogeneity in most (but not all) cell types (***Angelidis et al., 2019***), based on the distance-to-mean method of ***Enge et al***.. Later, a study on murine lung, spleen and kidney corroborated by Euclidean distance-to-centroid methods an age-related increase in cell-to-cell variability, albeit present in some cell types only (***Kimmel et al., 2019***). In contrast, a study in murine aging brain found no increase in transcriptional heterogeneity associated with aging (***Ximerakis et al., 2019***). Overall, these results suggest that the purported age-associated increase in transcriptional noise might be restricted to particular cell types or tissues. Of note, alternative explanations to the variability in the expression of individual genes being the basis for increased transcriptional noise do exist. Among others, the lack of balance between spliced and unspliced mRNAs (***Gupta et al., 2021***) and the existence of dysregulated gene regulatory networks (***Mishra et al., 2021***) have been proposed. In fact, Bashan and colleagues developed a novel computational tool to measure age-related loss of gene-to-gene transcriptional coordination (what they called *Global Coordination Level* or GCL), and reported a GCL decrease in aging cells across diverse organisms and cell types, which was also associated with a high mutational load (***Levy et al., 2020***). In a nutshell, these authors suggested that the observations of age-associated increase in cell-to-cell variability were restricted to specific cell types and tissues, but not generalized. Instead, they proposed that transcriptional dysregulation occurs at the level of gene-to-gene coordination. Despite the numerous attempts at measuring transcriptional noise in aged tissues, several challenges remain: (i) there are important differences in between studies with regard to the definition of *transcriptional noise* and the computational methods used to quantify it; (ii) studies focused mostly on single datasets of different tissues and cell types, while it is well known that both the inter-tissue and the inter-cellular variability might be significant; and (iii) little to no attention was given to the fact that cellular composition of aged organs shows relevant variability as compared to the young (***Nalapareddy et al., 2022***).

In the present work, we aimed to systematically measure age-associated transcriptional noise across different tissues and species, testing diverse computational methods in parallel. The main goals of the study were to substantiate claims of age-associated transcriptional noise increase and determine whether it presents a cell type-specific pattern. For this, we took advantage of the large number of aging mouse and human scRNAseq datasets that are publicly available, and developed two computational tools (*Decibel* and *Scallop*) to analyze them by focusing on two aspects: age-related transcriptional noise and changes in cell type composition.

## Results

### *Decibel*: a Python toolkit for transcriptional noise quantification

We developed a Python toolkit for the quantification of transcriptional noise in scRNAseq, where we implemented the four main families of methods that have been used in the literature to measure loss of cell type identity associated with aging (Figure 1A). The first method, which we refer to as *biological variation over technical variation*, takes the Pearson’s correlation distance between each cell and the mean expression vector of its corresponding cell type for that individual, using the whole transcriptome in the calculation. It then divides this correlation by the ERCC-based distance between each cell and its cell type mean. This method can only be used when ERCC spike-ins have been included in the experimental design. The second method computes the gene-based Euclidean distance between each cell’s expression vector and its cell type mean expression vector per donor/individual. The third one computes the Euclidean distance between each cell and the average gene expression across cell types, using a set of invariant genes. Invariant genes are selected by splitting the whole transcriptome into ten equally-sized bins according to their mean expression. Then, the two bins with the most extreme expression values are discarded and the 10% with the highest coefficient of variation within each of the remaining bins are selected. The fourth one is the GCL. Its original formulation takes a dataset containing a single cell type and it randomly splits its transcriptome into two halves, then computes the dependency between them as the batch-corrected distance correlation (***Levy et al., 2020***). The GCL is obtained by averaging this dependency over *k* iterations. We implemented an extension of this method so that it could be used in datasets containing several cell types, by computing the GCL averaged over 50 iterations for each cell type of the same individual. Therefore, our implementation outputs a GCL score per cell type and individual rather than a transcriptional variability measure per cell. The Python implementation of these four methods is available at https://gitlab.com/olgaibanez/decibel.

**Figure 1.**
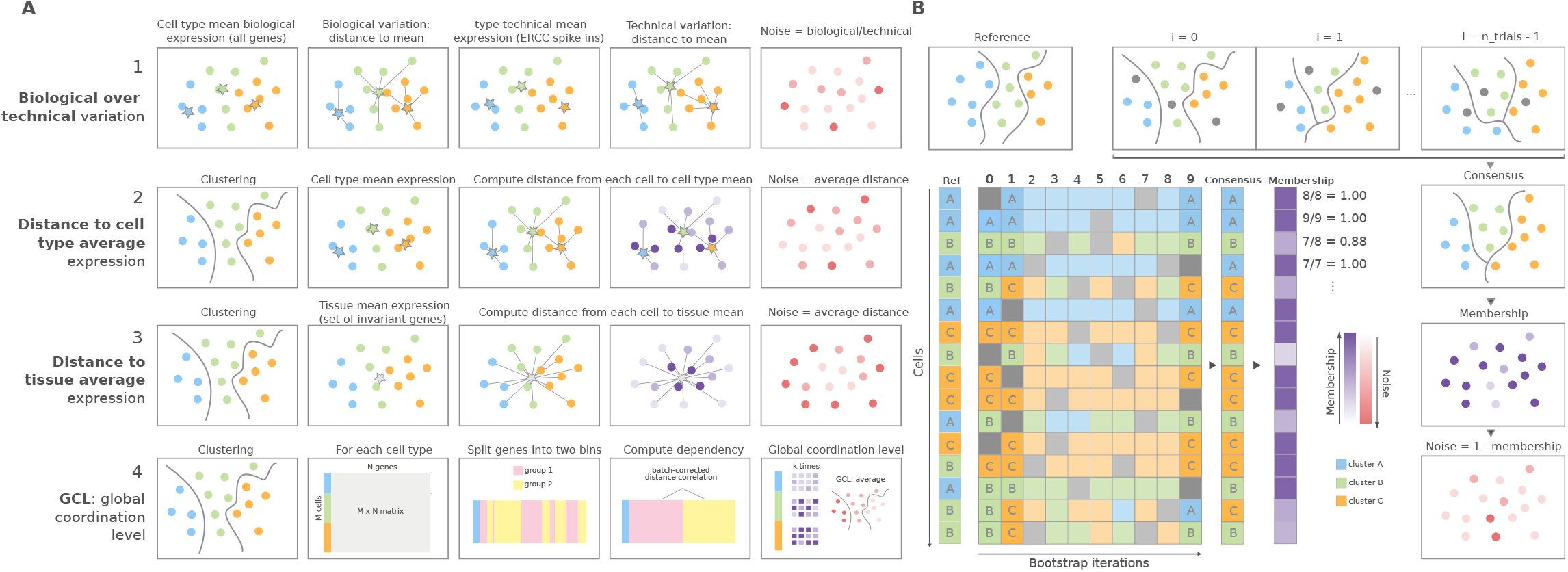
Overview of computational methods for the quantification of transcriptional noise and example workflow in *Scallop*. (A) The methods implemented in *Decibel* Python toolkit are summarized through diagrams depicting how they measure transcriptional noise. 1) Biological variation (whole transcriptome-based Pearson’s correlation distance between each cell and the mean expression vector), divided by the technical variation (ERCC spike-in based distance) (***Enge et al., 2017***) 2) Mean whole transcriptome-based Euclidean distance to cell type average (***Enge et al., 2017***) 3) Mean invariant gene-based Euclidean distance to tissue average. (***Enge et al., 2017***) 4) GCL (***Levy et al., 2020***) per cell type. Stars represent the “center” of each cluster (average gene expression for each cell type). (B) *Scallop*: example workflow on a 16-cell dataset. A reference clustering solution (*Ref*) is obtained by running a community detection algorithm (default: Leiden) on the whole dataset. Three clusters are obtained: A (blue), B (green) and C (orange). Then, a subset of cells is randomly selected and subjected to unsupervised clustering *n_trials=10* times (cells not selected in each bootstrap iteration are shown in grey). The cluster labels across bootstrap iterations are harmonized by mapping the cluster labels with the greatest overlap, using the Hungarian method (***Munkres, 1957***). A consensus clustering solution is derived by selecting the most frequently-assigned cluster label per cell, and the membership score is computed as the frequency with which the consensus label was assigned to each cell. *Scallop* measures noise as a 1 - membership value assigned to each cell. **Figure 1–Figure supplement 1. Performance of *Scallop* in comparison to preexisting methods for the quantification of transcriptional noise**. The different methods were tested on a dataset of 8,278 human T lymphocytes. (A) UMAPs and dotplot showing *CD3, CD4* and *CD8* marker gene expression per cluster. (B) Representation of transcriptional noise levels, as measured by using two distance-to-centroid methods (*euc_dist* and *euc_dist_tissue_invar*), 1 - membership (*scallop_noise*) and Global Coordination Level (*GCL*). (C) The 10% most stable (purple) and 10% most unstable (red) cells are represented on the UMAP plots for *Euclidean distance to cell type mean* (top row) and *Scallop* methods (bottom row), respectively. **Figure 1–Figure supplement 2. *Scallop* robustness in relation to input parameters**. The plots on the left show the median correlation distance between membership scores of different runs of *Scallop* against (A) the number of trials, (B) the fraction of cells used in each bootstrap and (C) the resolution given to the clustering method (Leiden) in five independent scRNAseq datasets (PBMC3K, ***Joost et al. (2016***); ***Paul et al. (2015***); ***Moignard et al. (2015***), Heart10K). The median correlation distance was computed over 100 runs of *Scallop*. The swarmplots on the right show the distribution of the correlation distances between membership scores against each of the input parameters for the heart10k dataset. The median is shown as a red point. While, for the sake of clarity, a random sample of 100 correlation distances is shown for each value of the parameter under study, the median was computed using all the correlation distances. *Scallop* membership scores converge as we increase the number of bootstrap iterations and the fraction of cells used in the clustering. **Figure 1–Figure supplement 3. Stable cells as identified with *Scallop* are more representative of the cell type than unstable cells**. Distribution of log-fold changes (top row) and adjusted *p*-values (middle row) of the first 100 differentially expressed genes (DEGs) between each cell type or subtype and the rest of the cells in six cell types and subtypes from the 10X PBMC3K dataset. The overlap between the DEGs found when using all of the cells, only the stable cells and only the unstable cells is also shown (bottom row). The adjusted *p*-values obtained with all the cells are equivalent to those obtained using only the most stable half of the cells. In contrast, the differential expression of many genes is not statistically significant when using the unstable half from each population. The overlap between the top 100 DEGs obtained is very high between the stable cells and all cells subsets, whereas DEGs obtained in unstable cells have a very low intersection with all cells.

### *Scallop* membership score accurately identifies transcriptionally noisy cells

In addition to implementing existing methods, we developed *Scallop*, a novel tool for the quantification of the degree of loss of cell type identity in scRNAseq data (Figure 1B). *Scallop* measures the membership of each cell to a particular cluster by iteratively running a clustering algorithm on randomly selected subsets of cells and computing the fraction of iterations a cell was assigned to a particular cluster. Thus, cluster membership takes values between 0 and 1. *Scallop* relies on the assumption that the more consistently a cell is assigned to a particular cluster across bootstrap iterations, the greater its transcriptional stability. Conversely, a cell being assigned to different clusters across iterations is indicative of a greater transcriptional variation. Therefore, we quantify loss of cell type identity as 1 − membership.

In order to characterize and validate the performance of our method, we conducted three experiments. First, we compared the output of *Scallop* to the transcriptional noise measured using the methods implemented in *Decibel* on 8,278 human T cells drawn from the Peripheral Blood Mononuclear Cell (PBMC) 20K dataset from 10x Genomics. Clustering revealed three main T cell subtypes, which we annotated according to their expression of *CD4* and *CD8* markers (Figure 1 - Supplement 1A). Then, we measured transcriptional variability and gene coordination level using *Decibel* and inspected the distribution of variability scores over the Uniform Manifold Approximation and Projection (UMAP) plots (Figure 1 - Supplement 1B). Unlike distance-to-centroid methods, *Scallop* detected transcriptionally noisy cells that lie in between transcriptionally stable T cell subtypes on the UMAP plot. GCL yielded different coordination levels for each T cell subtype, but the method does not allow for comparisons between individual cells, as it outputs a single score per cell type.

In addition, we plotted the 10% most transcriptionally stable and unstable cells according to the *Euclidean distance to the cell type mean* and *Scallop* methods (Figure 1 - Supplement 1C). These analyses suggested that *Scallop*’s membership score outperforms distance-to-centroid methods at discriminating between noisy cells lying in between clusters and more transcriptionally robust cells constituting the core of T cell subtypes.

Next, we analyzed *Scallop*’s robustness in response to input parameters, namely, the number of bootstrap iterations and the fraction of cells used in each iteration. We ran *Scallop* on five independent scRNAseq datasets with different size and cluster composition (see Appendix 1), and studied the convergence of *Scallop* membership scores for a wide range of values (Figure 1 - Supplement 2). The median correlation distance between membership scores decreased as we increased the number of bootstrap iterations (`n_trials`) and the fraction of cells used in each iteration (frac_cells). We concluded that *Scallop*’s output is robust to changes in its input parameter values, the results suggesting that frac_cells > 0.8 and n_trials > 30 are appropriate parameter values for most datasets (Figure 1 - Supplement 2).

Finally, we studied the relationship between *Scallop* membership score and robust gene marker expression, by comparing the statistical significance of the output of differential expression analysis between cell type clusters, conducted on stable and unstable cells. For this, we analyzed six cell types and subtypes (*CD4* and *CD8* T cells, *CD14* and *FCGR3A* monocytes, dendritic cells, and NK cells) from the PBMC 3K dataset from 10x Genomics. Cells with a higher *Scallop* membership to their cluster differentially expressed cell type-specific markers with greater statistical significance than low-membership cells (Figure 1 - Supplement 3). Overall, these results showed that *Scallop* membership is related to a more robust expression of gene markers defining cell types than other existing methods.

### Increased transcriptional noise is not a universal hallmark of aging

To determine if aging is associated with a generalized increase in transcriptional noise at the tissue level, we used *Scallop* to compare the average degree of membership of young and old cells to their cell type cluster in scRNAseq datasets of various tissues (Figure 2). For the initial analysis, we selected seven datasets where transcriptional noise had already been measured using different methods, and with differing outcomes (Appendix 1 summarizes the main characteristics of each dataset as well the findings regarding transcriptional noise obtained in each of the studies). The age and cell type composition of the final datasets used in our study are shown in Figure 2 - Supplement 1, and the samples included in the datasets as well as the inclusion criteria are provided in Appendix Additionally, the methods implemented in *Decibel* to compute loss of identity were run in parallel as a control (Figure 2 - Supplement 2). When measuring the *Scallop* membership score of individual young and old cells to their cell type clusters, the results were inconsistent. Differences between age groups were found in some datasets, but the directionality of the change was not conserved across datasets, neither in the average 1 − membership score, nor in the percentage of noisy cells in the young and the old fraction of each dataset. For most datasets (***Ximerakis et al., Angelidis et al., Martinez-Jimenez et al., Kimmel et al***.), no significant change in mean transcriptional noise was found. Two datasets (***Enge et al***. and ***Salzer et al***.) showed an increase mean membership associated with aging, although we observed the interquartile range of noise values to be very similar between young and old individuals. In one of the datasets (***Solé-Boldo et al***.), cells showed decreased transcriptional noise with aging. Of note, similar inconsistent results were found when using the preexisting noise-measuring methods as compiled in *Decibel*, even when applying different methods to the exact same dataset (Figure 2 - Supplement 2). Overall, these results indicated that a generalized *increase in transcriptional noise* or a *loss of cellular identity* are not universal hallmarks of aging, at least at the tissue level. However, the possibility that transcriptional noise increased in specific cell types was still unexplored by these analyses.

**Figure 2.**
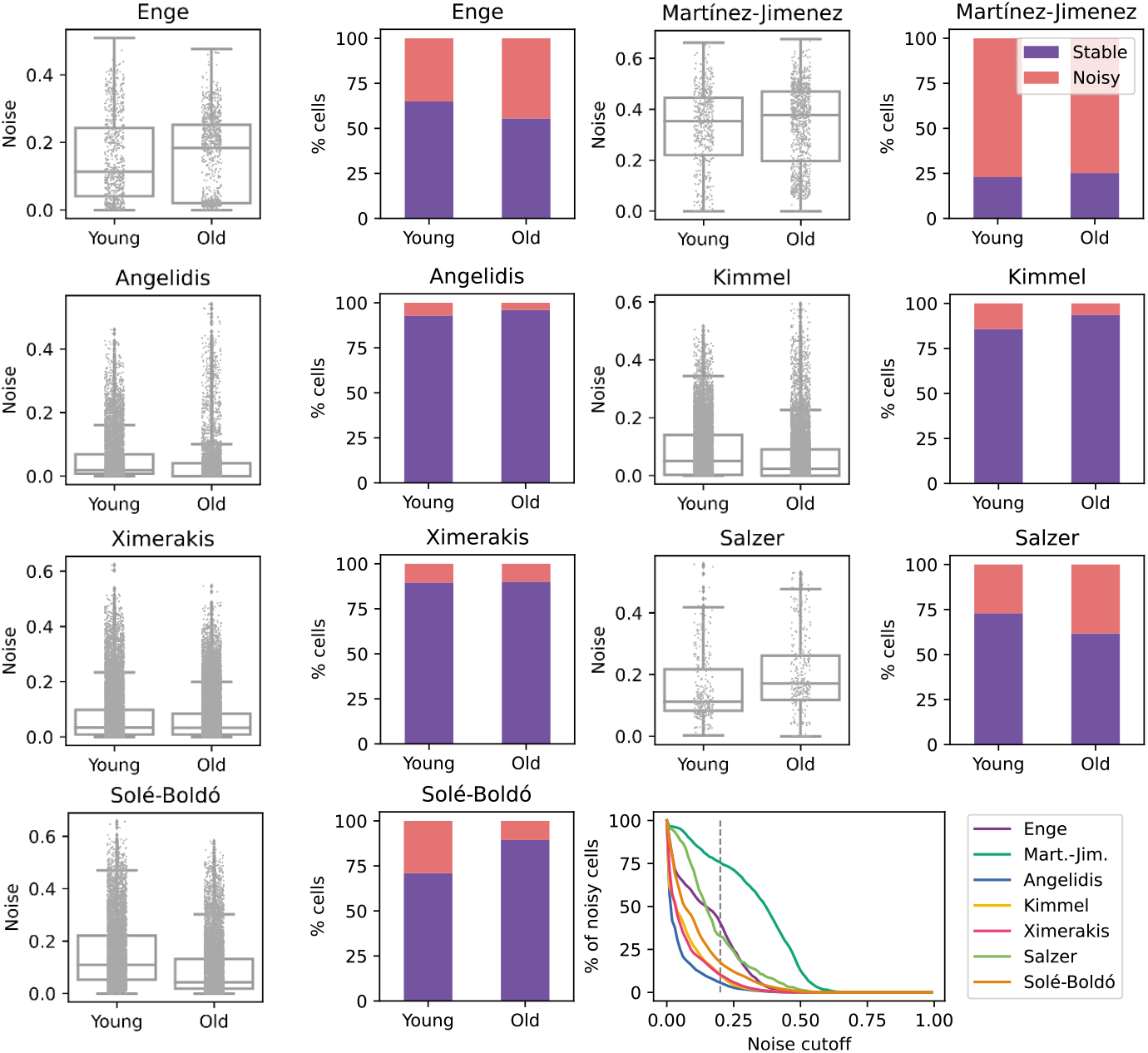
No consistent increase in transcriptional noise detected in seven scRNASeq datasets of aging at the tissue level. The graphs show the amount of transcriptional noise, computed as 1 - *membership* to cell type clusters, in the young and old age groups of seven single-cell RNAseq datasets of different tissues. For each dataset, the distribution of transcriptional noise values is shown as a stripplot over a boxplot, where the whiskers represent 1.5 times the interquartile range. Next to them the proportions of stable and noisy cells (noise ≥0.2) per age group are shown (Purple bars=stable cells, pink bars=noisy cells). At the bottom right panel, the percentage of noisy cells with a transcriptional noise over a cutoff of 0.2 is plotted against the cutoff. Each colored line represents a different dataset. **Figure 2–Figure supplement 1. Composition of the seven scRNAseq datasets of aging used in this figure**. UMAPs showing the age and cell type composition of the seven datasets used in the analysis of the age-related transcriptional noise at the tissue level. The UMAPs show the final composition of the datasets used in the experiment. The cell type annotations were obtained from the original authors in all datasets except Kimmel lung, where the labels from Angelidis were projected onto the dataset. **Figure 2–Figure supplement 2. Measurements of transcriptional noise on seven scRNAseq datasets of aging using computational methods implemented in *Decibel***. Stripplots showing the distribution of noise values, as measured by the four alternative methods (Biological variation over technical variation, Euclidean distance to cell type mean, Euclidean distance to tissue mean using invariant genes, and Global Coordination Level - GCL) in the seven datasets used in the analysis of the age-related transcriptional noise at the tissue level. Boxplots and their whiskers represent the interquartile range (IR) and 1.5*IR respectively.

### The murine aging lung shows no consistent pattern of transcriptional noise at the cell type level, and is instead characterized by reproducible alterations in immune cell composition

Do specific cell types become noisier as they age? In order to answer this question, we focused on a single tissue and conducted an in-depth analysis of transcriptional noise at the cell type level. For this, we selected the murine aging lung because of the relative abundance of available datasets in which authors had reported an age-associated increase in transcriptional noise, yet restricted to particular cell types: ***Angelidis et al., Kimmel et al***.; and the Tabula Muris Senis (TMS) lung droplet and FACS datasets (***Almanzar et al., 2020***). In each dataset, transcriptional noise was measured as 1 − membership to cell type clusters in the young and old fractions, and the differences in median noise between the old and the young fraction for each of the existing 31 lung cell types and subtypes were calculated (Figure 3). Since changes in the gene expression of tissues can also be caused by altered cell type composition (***Trapnell, 2015***), we estimated the relative abundances of the 31 cell types in the young and old fraction of each dataset and measured the effect of age by fitting Generalized Linear Models (GLM) to cell type composition data of the four datasets, using each mouse as a biological replicate (Figure 3 - Supplement 1). By plotting the age-related cell type enrichment together with the cell-to-cell transcriptional variability in each of the datasets, we obtained a comprehensive map of cell type enrichment and transcriptional noise associated with aging at the cell-identity level (Figure 3A). In this analysis, the direction and magnitude of changes in transcriptional noise varied across cell types. For instance, club cells (a bronchiolar exocrine cell type) were detected in sufficient numbers in 3 out of 4 datasets, their median membership score consistently decreasing 10-17% (which showed up as a moderate increase in transcriptional noise in Figure 3A; bubble #22). Similarly, lung interstitial fibroblasts’ transcriptional noise appeared to increase with age, although with a larger range of membership scores (3-17%; bubble #24). In both cases, the cell type abundance was not affected by aging. In contrast, alveolar macrophages showed a decrease in age-associated transcriptional noise (5-12% increase in median membership; bubble #10). Finally, several cell identities appeared not to change significantly with regard to their transcriptional noise related to aging. That was clearly the case for capillary endothelial cells (bubble #9) and plasma cells (bubble #5). Vascular endothelial cells (bubble #6) showed less than 2% of change in noise in 3 out of 4 datasets, but increased up to 8% in one dataset. Therefore, and contrary to expectation, quantitative analysis of age-associated transcriptional noise did not show a consistent pattern across diverse lung cell types in the four available datasets.

**Figure 3.**
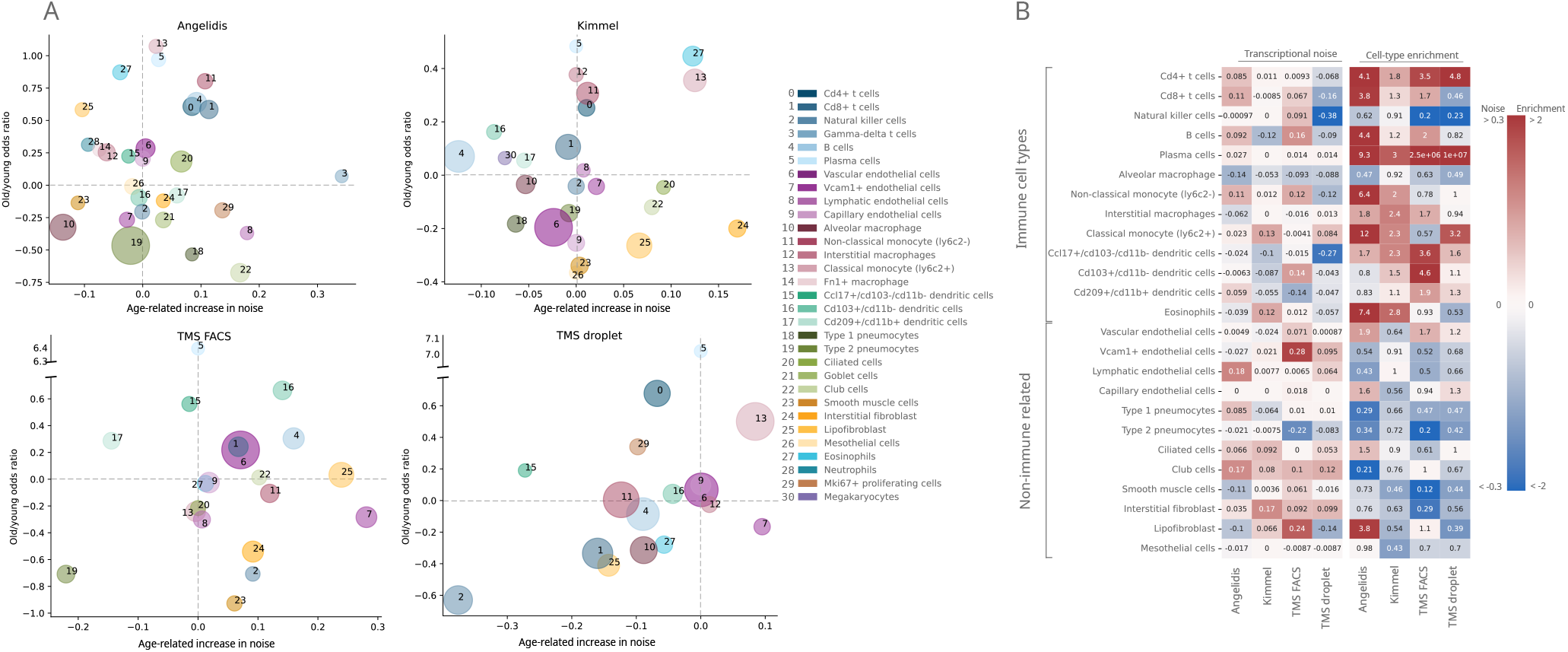
Lack of evidence for an increase in transcriptional noise of the murine aging lung and detection of an enrichment in immune cells. (A) Bubble chart of transcriptional noise and cell type enrichment (old/young OR) of 31 murine lung cell identities. The age-related change in transcriptional noise (horizontal axis) is computed by *Scallop* as the decrease in median membership score per cell identity between young and old cells. The enrichment of each cell type in old samples with respect to their young counterpart is represented as the old/young OR (vertical axis). The area of the bubbles represents the expected proportion of each cell type in the whole dataset according to the binomial GLM fitted for that dataset. (B) Immune cell type enrichment, but not age-associated increase in transcriptional noise, is consistently detected in old mice lungs. The increase in transcriptional noise associated with aging (left) and the cell type enrichment (right) are shown for several lung cell identities classified on the left as immune and non-immune. Cell identities present in at least 3 out of the 4 studied datasets are shown, and the age-related difference in transcriptional noise of missing cell identities is imputed from the remaining three measurements (mean difference across datasets). **Figure 3–Figure supplement 1. Composition of the four scRNAseq datasets of the murine aging lung used in this figure**. (a) Experimental approach. Four murine aging lung datasets were preprocessed and cell type-annotated. The cell-type labels from Angelidis were used as a reference to annotate the rest of the datasets. Differences in cell-type abundance between young and old mice were quantified using GLMs. From each dataset, eight subsets of related cell-types were created to classify the 31 cell types into 8 categories, which were used as input for *Scallop* to analyze the differences in cell-to-cell variability. (b) Cell type-annotated mouse lung datasets. UMAP plots showing the four datasets with their cell type annotations. **Figure 3–Figure supplement 2. Qualitative ranking of murine aging lung cell types according to transcriptional noise and cell type enrichment**. The 31 detected lung cell types were classified in the *Noise* ranking (left) according to their greater age-related increase in noise. They were also classified in the *Enrichment* ranking (right) according to their greater enrichment in old samples. Cell categories that were represented by fewer than 100 cells were excluded from the transcriptional noise evaluation, and therefore do not appear in the plot. Specific cell types are shown in the same color and with the same numbers as specified in the legend. **Figure 3–Figure supplement 3. Comparison of the originally reported cell type-associated increase in transcriptional noise with the results obtained with *Scallop***. The content of the first three columns was drawn from the original publications (***Angelidis et al***.; ***Kimmel et al***.). More specifically, *Angelidis_TN* is the transcriptional noise per cell identity on the Angelidis dataset (from their figure 2); *Kimmel_OD* is the gene overdispersion per cell type on the Kimmel dataset (from their figure 2B); and *Kimmel_DC* is the cell-cell heterogeneity per cell identity measured as the Euclidean distance to the centroid of the cell identity for a particular age. Columns 4-7 summarize the results of our analysis of age-related loss of cell type identity in the murine lung. Specifically, *Angelidis_S, Kimmel_S, TMS_FACS_S and TMS_drop_S* report the transcriptional noise per cell identity on the four datasets, measured as the difference in median membership score between young and old individuals. The cell identities used are those drawn from Angelidis. Since some cell identities from Kimmel dataset did not have a 1:1 correspondence to the Angelidis cell identities, they are shown using their original notation at the bottom of the table (“Additional cell identities”). UP/DOWN: age-related increase/decrease in noise, NS: the difference in noise between young and old individuals is not statistically significant. NP: the cell identity was not present in the dataset in sufficient amounts to perform the analysis. For most cell types, it can be concluded that there is little overlap between cell identity-specific noise measurements across datasets and methods

In contrast, the cell abundance analysis did reveal a strikingly consistent enrichment of immune cell types (lymphocytes in particular) across all datasets in old samples, indicative of immune cell infiltration in the old tissue. In particular, plasma cells (bubble #5) showed highly consistent enrichment in old mice, with an Old/Young odds ratio (OR) of 3 in the Kimmel dataset (*p*-value=1.1 × 10−5) and of 9.3 in the Angelidis dataset (*p*-value=6.5 × 10−21). The ORs for the two TMS datasets were most likely overestimated due to low cell numbers (only 9 and 22 old plasma cells were detected in the TMS datasets). The more abundant B cells (bubble #4) were also significantly enriched in 3/4 datasets (Angelidis: OR=4.4, *p*-value=2.5 × 10−69; Kimmel: OR=1.2, *p*-value=6.3 × 10−8; TMS FACS: OR=2.0, *p*-value=8.9 × 10−6). Other immune cell types such as monocytes, macrophages and dendritic cells also appeared to be enriched in all datasets. This prompted us to further investigate the basis for the apparent immune cell enrichment and its potential relationship to increased transcriptional heterogeneity in the old age. In a qualitative approach to look for consistent patterns across datasets, we ranked cell identities according to their age-related increase in noise and enrichment (Figure 3 - Supplement 2). While most cell types were evenly distributed along the transcriptional noise ranking, this representation provided a visible distinction between immune and non-immune cell types regarding their age-related enrichment, with nearly all immune cell types appearing on top of the enrichment ranking. For instance, plasma cells (Figure 3 - Supplement 2, #5) were the third most enriched cell type in the Angelidis dataset and appeared on the top position in the rest of the datasets. Classical monocytes (#13) were found within the top 4 most enriched in 3/4 datasets. Interestingly, natural killer (NK) cells (#2) were the only underrepresented lymphocytes in old mice, and ranked consistently in the least enriched positions among immune cells. Conversely, parenchymal cell types such as goblet cells, club cells and ciliated cells consistently appeared at the bottom of the enrichment ranking, indicating that their proportion diminished with increased immune cell infiltration in the organ or, alternatively, loss of parenchymal cells associated with the old age. Endothelial cells were more evenly distributed along the ranking and thus did not show a clearly discernible age-associated enrichment or loss. Finally, we separated the lung cells into immune and non-immune cell categories and represented transcriptional noise and cell type enrichment values on a heatmap (Figure 3B). As clearly seen in this representation, the transcriptional noise increase associated with aging was extremely variable across cell identities and not always consistent across datasets (a comparison between these results and the results reported by ***Angelidis et al***. and ***Kimmel et al***. is provided in Figure 3 – 3). In contrast, the immune vs non-immune cell distinction alone explained the behaviour of most cells with respect to their relative abundance with very few exceptions, namely NK cells and alveolar macrophages.

### Changes in the abundance of the immune and endothelial cell repertoires characterize the human aging lung

Our analysis of age-related cell type enrichment and increase in transcriptional noise in the murine lung highlighted the importance of the changes associated with the relative abundance of cell types that conform the aging lung. To test if this was specific of murine lungs or it could be a more generalized phenomenon, we conducted similar analyses on two large scRNAseq datasets of the aging human lungs (15,852 cells from nine donors from the ***Raredon et al***. dataset and 15,048 cells from two donors of the Human Lung Cell Atlas (HLCA) dataset by ***Travaglini et al***.; (***Raredon et al., 2019; Travaglini et al., 2020***)). We harmonized cell type labels between datasets by projecting the HLCA labels onto the Raredon dataset (Figure 4 - Supplement 1). Then, we calculated the difference in mean membership score between old and young cells for each cell type in the two datasets, together with the cell type enrichment using the GLM method as described earlier (Figure 4). In general, and similar to what we had previously observed in the murine aging lung, we found a lack of consistency between the two datasets regarding transcriptional noise associated with aging of specific lung cell types. However, we did observe some conserved changes in cell type composition. Particularly, many immune cell types were enriched in older donors, as in the murine aging lung. Plasma cells were significantly enriched in the Raredon dataset (OR=2.6, *p*- values=1.9e-6) and enriched, albeit not significantly, in the HLCA dataset (OR=3.5e-11, *p*-value=1). The latter result was most probably due to lack of statistical power, as the dataset only consists of two donors. Interestingly, alveolar macrophages were enriched (rather than depleted as in the murine aging lung) in both human aging lung datasets (OR=1.2, *p*-value=3.61e-5 in Raredon; OR=2.8, *p*-value=1.54e-261 in HLCA). Several endothelial cell types were significantly depleted in the two human aging lung datasets. Vein endothelial cells (OR=0.65, *p*-value=7.7e-5 in Raredon; OR=0.58, *p*-value=1.9e-9 in HLCA), capillary endothelial cells (OR=0.81, *p*-value=3.4e-2 in Raredon; OR=0.3, *p*-value=3.5e-141 in HLCA), endothelial cells of lymphatic vessels (OR=0.51, *p*-values=1.9e-9 in HLCA). These results indicated that aged human lungs present reduced vascularization and significant immune cell infiltrates as compared to the young. These facts may have influenced previous analyses of transcriptional noise associated with aging.

**Figure 4.**
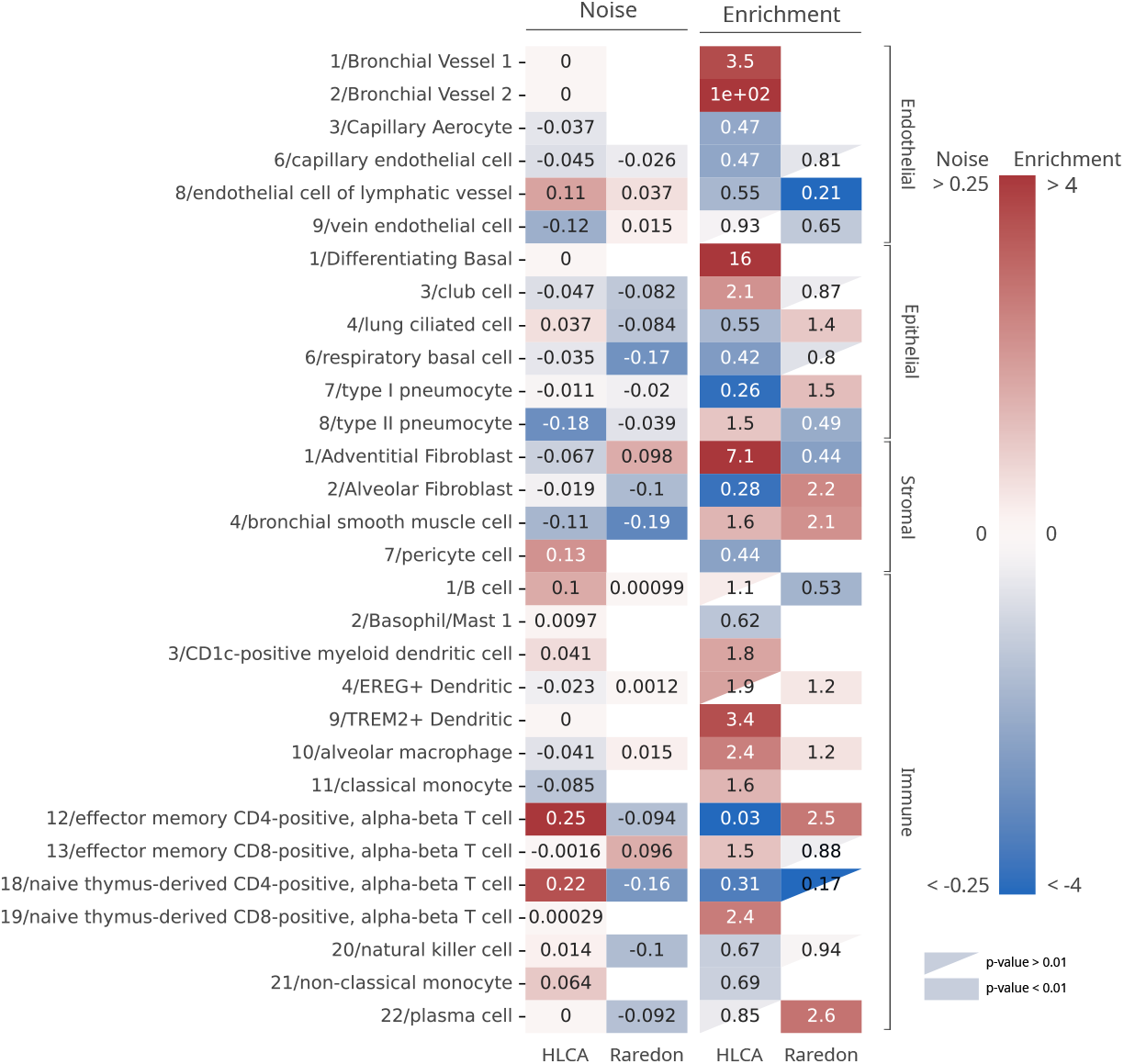
Human aging lungs show no increase in transcriptional noise but consistent depletion and enrichment of specific endothelial and immune cell types. The increase in transcriptional noise associated with aging (*Noise*, left) and the cell type enrichment (*Enrichment*, right) values are shown for 30 human lung cell identities as detected in the HLCA and Raredon datasets (***Raredon et al., 2019; Travaglini et al., 2020***). For each cell type, its age-related increase in noise (difference in 1 - *membership* between old and young individuals per cell type) and the Old/Young OR are shown. Only cell types whose enrichment/depletion are statistically significant in at least one of the datasets are shown, and the OR-s associated with a *p*-value > 0.01 are shown as a triangle. The color-bar for the enrichment is shown in a logarithmic scale. **Figure 4–Figure supplement 1. Composition of the two scRNAseq datasets of the human aging lung used in this figure**. The UMAP plots with the age and cell type identity annotations are shown for each tissue compartment (endothelial, epithelial, stromal and immune) and each dataset separately.

### Distance-to-centroid methods detect transcriptionally stable cell subtypes as transcriptional noise

A relevant open question is what was the source of apparent transcriptional *noise* in previous studies that were based on distance-to-centroid methods. Since we found important changes in the community of human alveolar macrophages in the HLCA dataset, we conducted an in depth analysis on that cell type that revealed four distinct alveolar macrophage communities that emerge with aging from a single transcriptionally homogeneous cluster (see Figure 5 and Supplement 1). The four aged alveolar macrophage subclusters present a markedly different expression of genes coding for surfactant proteins (*SFTPA1, SFTPA2, SFTPB, SFTPC*, and *SFTPD*), i.e. they show changes consistent with alternative fate determination. Deregulated surfactant protein expression is connected to the age-related functional decline of human lungs. In fact, mutations in the gene coding for surfactant protein C (*SFTPC*) and in the *MUC5B* promoter region are linked to pulmonary fibrosis, but the effects of these mutations are usually not observed until late in life (around 60-70 years old), because age-related decline in proteostasis is needed for aggregation prone or misfolded proteins to actually cause damage (***Schneider et al., 2021***). Interestingly, we measured the age-associated transcriptional noise in the alveolar macrophage community using a distance-to-centroid method (*Euclidean distance to the cell type mean*) and *Scallop*, and observed that only the latter algorithm could accurately detect the emergence of distinct and transcriptionally stable alveolar macrophage subpopulations, whereas the distance-to-centroid method would interpret this specification of alternative cell fates as a single macrophage population undergoing loss of identity (Figure 5).

**Figure 5.**
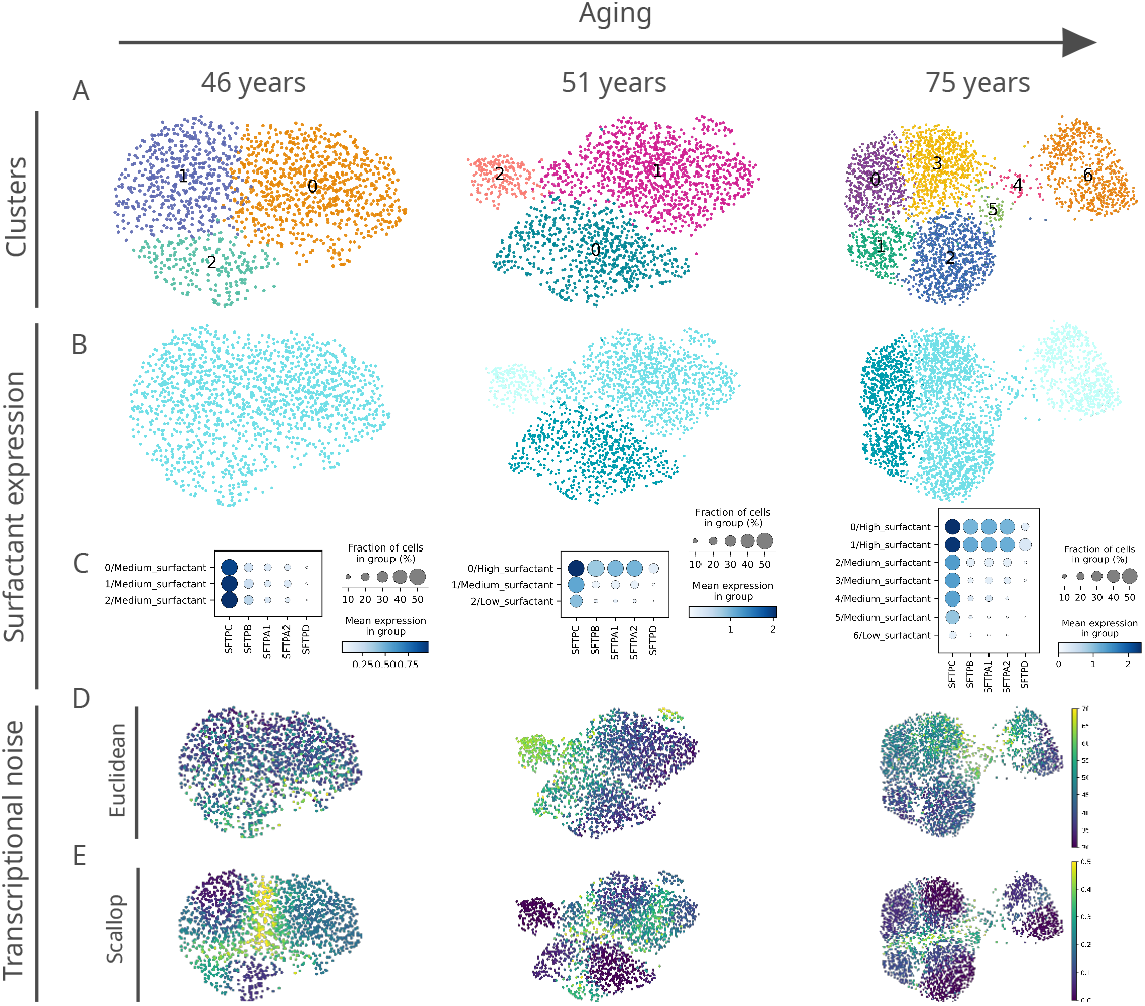
Euclidean distance-to-centroid methods are unable to distinguish *bona fide* transcriptional noise from alternative cell fate specification. (A) An increasing number of alveolar macrophage subclusters (as obtained with Leiden) are detected in three donors (aged 46, 51, and 75 years) from the ***Travaglini et al***. (HLCA) dataset. (B-C) The new cell clusters are characterized by differential surfactant protein gene expression levels, as clearly seen on the UMAP (B) and dotplot (C) representations. (D-E) Transcriptional noise measurements, using the Euclidean distance to cell type mean (D) and 1 - *membership* using *Scallop* (E), demonstrate that only the latter method is able to distinguish *bona fide* transcriptional noise from the formation of new clusters that are transcriptionally stable. **Figure 5–Figure supplement 1. Expression of surfactant protein genes by human alveolar macrophages**. Differential expression by alveolar macrophage cell clusters of the genes coding for surfactant proteins *SFTPA1, SFTPA2, SFTPB, SFTPC*, and *SFTPD* is shown for three donors (aged 46, 51 and 75) of the ***Travaglini et al***. (HLCA) dataset.

## Discussion

Mechanistically, it is not clear if aging is a tightly regulated process or may be the result of passive phenomena of stochastic nature (***Schmeer et al., 2019; Gladyshev, 2016; Costa et al., 2016***). In the absence of further mechanistic insight, aging is characterized by a series of phenotypic changes at the cellular and tissue levels, such as genomic instability, epigenetic alterations, chronic low level inflammation (*inflammaging*), immunosenescence, and impaired regeneration (***López-Otín et al., 2013; Gems and de Magalhães, 2021***). In addition, an increase in transcriptional noise has been observed in some aged tissues and cell types (***Enge et al., 2017; Martinez-Jimenez et al., 2019; Angelidis et al., 2019; Kimmel et al., 2019***). Transcriptional noise could be related to genomic instability (***Vijg, 2021***), epigenetic deregulation (***Lu et al., 2020b; Oliviero et al., 2022***) or loss of proteostasis (***Li et al., 2020***), all established hallmarks of aging. Some authors consider transcriptional noise to be a hallmark of aging in and of itself (***Mendenhall et al., 2021***). Since aging is multifactorial and mutational load most likely leads to clonal expansion of aberrant cells that accumulate throughout the lifetime of the individual, other authors suggest that aging traits may be associated with cell type imbalance in aged organs (***Cagan et al., 2022***). Another recent hypothesis is *inter-tissue convergence* through age-associated loss of specialization (***Izgi et al., 2022***).

In this work, we made a systematic comparison of the most important families of methods that have been used to quantify age-related transcriptional noise through the implementation of *Decibel*, a novel Python toolkit. Since we were not convinced of the utility of these methods to determine *bona fide* transcriptional noise, we developed a novel method and applied it to a wide array of tissues. Our proposed tool, *Scallop*, presents some advantages over existing methods: it does not require neither ERCC spike-ins nor cell type labels. In addition, it provides information that is complementary to the GCL, as it yields a cell-wise measurement of transcriptional noise that enables us to compare between *stable* and *unstable* cells within the same cluster or cell type. Most importantly, *Scallop* measures transcriptional noise by membership to cell type-specific clusters which is a re-definition of the original formulation of *noise* by ***Raser and O’Shea***. This is in stark contrast to measurements of *noise* including other phenomena (as demonstrated in Figure 5) by the distance-to-centroid methods prevalent in the literature. When applied to seven independent aging datasets, the results obtained revealed little overlap in the magnitude and directionality of the changes in transcriptional noise associated with aging of the different tissues analyzed, providing evidence that an increase in transcriptional noise might not be as evident as generally thought.

In order to investigate cell type-specific effects in transcriptional noise, it is crucial to compare between different datasets of the same aging tissue. Otherwise, it is difficult to ascertain whether the variability observed between cell types is due to a pattern that is conserved in that tissue or is merely the effect of the intrinsic variability associated with scRNAseq experiments (***Fonseca Costa et al., 2021***). For the cell type-specific study, we focused on the aging lung, as the effect of aging of this tissue has has gained relevance (***Schiller et al., 2019***) due to its association with chronic obstructive pulmonary disease (COPD), lung cancer and interstitial lung disease (***Angelidis et al., 2019; Schneider et al., 2021***), and its increased risk of severe illness in COVID-19 patients (***Williamson et al., 2020***). In the 31 cell types analyzed in mouse lungs, we found increased transcriptional noise in club cells and interstitial fibroblasts only, while alveolar macrophages seemed to decrease it. Of interest, a single-cell analysis of alveolar macrophages did not identify distinct clusters associated with mouse or human aging, identifying changes in the aged alveolar microenvironment as key for their altered functionalities (***McQuattie-Pimentel et al., 2021***). In humans, we analyzed two aging lung datasets that provide complementary information, as the final Raredon dataset consists of 9 donors of a wider range of ages but is not as well-powered in terms of cell type resolution as the HLCA dataset, which contains 48 cell identities. Similar to what we had previously observed in the murine aging lung, there was no consistency between the two datasets regarding transcriptional noise of the 30 specific cell types detected. However, both in human and mouse lungs we detected a shift in the abundance of a number of cell populations with age, most clearly seen for immune cells.

In fact, the age-associated increase in immune cell infiltration of solid organs may be generalized. Specifically, one study found neutrophil and plasma cell infiltration in adipose tissue, aorta, liver and kidneys of aged rats of both sexes, and the immune cell infiltration was reversed by caloric restriction (***Ma et al., 2020a***). Another study found a subtype of highly secretory plasma cells infiltrated in the aged bone marrow, spleen, fat, kidney, heart, liver, muscle, and lungs (***Schaum et al., 2020***). Of note, immune cell senescence has been shown to induce aging of solid organs (***Yousefzadeh et al., 2021***), in what has been proposed to be a feed-forward circuit (***Salminen, 2021***). Therefore, the importance of immune cell infiltration of the aged lungs cannot be overlooked. In fact, age-associated immune cell type enrichment has also been observed in two independent studies of macaque lungs. One study found increased mast cells, plasma cells and CD8+ T cells in aged lung tissue (***Ma et al., 2020b***), while the other found increased alveolar and interstitial macrophage numbers in bronchoalveolar lavages of old macaques (***Rhoades et al., 2022***). The significance of the shift in cellular composition of the aged lungs in relation to the appearance of aging traits remains to be determined. Of note, alternative explanations for transcriptional changes associated with aging such as *tissue convergence* are compatible with shifts in the cellular composition of aging tissues and organs being a primary cause of convergence (***Izgi et al., 2022***).

In summary, the sources of the apparent increase in *transcriptional noise* reported by previous studies may be multiple, and are mostly related to the computational methods used to characterize transcriptional noise and cellular identity in aged tissues. Open availability of *Decibel* and *Scallop* represents an opportunity for the aging research community to further investigate these issues, and they are also valuable for researchers addressing cell-to-cell variability of scRNAseq datasets in other settings.

## Methods

### *Decibel*: Python toolkit for the quantification of transcriptional noise

We developed a Python toolkit for the quantification of loss of cell type identity associated with aging. We implemented methods as they were originally described in the literature.

#### Biological variation over technical variation

Biological variation over technical variation is measured as in the original formulation by ***Enge et al***., by computing 1 − *ρ*, where *ρ* is the Pearson’s correlation between the gene expression vector of each cell and the mean expression of its cell type, i.e., the gene expression averaged over all the cells from the same cell type and individual. For each cell type and individual mouse or donor, the mean gene expression vector – the averaged expression of the whole set of monitored genes across cells – is computed. Then, the biological variation is measured as the Euclidean distance from each cell to its cell type mean for that individual. The technical variation is computed using the same procedure, but using only the ERCC spike-ins in the calculation of the distance to cell type mean. Finally, the transcriptional noise is calculated by dividing the biological variation by the technical variation per cell.

#### Euclidean distance to cell type mean

The distance to the cell type mean is measured as the second method described by ***Enge et al***.. For each cell type and individual mouse or donor, we compute the average whole-transcriptome expression. The noise is quantified as the Euclidean distance between the gene expression vector of each cell and its corresponding individual-matched cell type mean expression vector.

#### Invariant gene-based Euclidean distance to tissue mean

This is the third method described by Enge et al. (***Enge et al., 2017***). It is computed as the Euclidean distance from each cell to the average expression across cell types using a pre-selected set of invariant genes that is selected as follows: first, genes are sorted according to their mean expression and split into 10 equally sized bins, and the two extreme bins are discarded (10% most expressed and 10% least expressed genes). Then, the 10% of genes with the lowest coefficient of variation within each bin are selected and used for the calculation of the Euclidean distance between the mean expression vector across cell types and each of the cell expression vectors.

#### Average Global Coordination Level

Taking the Matlab code provided by the authors, we implemented the GCL in Python. As the original formulation was used in datasets with a single cell type, here we computed the GCL for each cell type separately and then calculated the average GCL for the tissue. For each cell type, the GCL was calculated by splitting the whole transcriptome into two random halves and computing the batch-corrected distance correlation between them (***Levy et al., 2020***). The GCL per cell type was averaged over k times. Following the authors’ recommendation, we used k=50 in all of our calculations.

### Scallop

*Scallop* iteratively runs a clustering algorithm of choice on randomly selected subsets of cells. Then, it computes the frequency with which each cell is assigned to the most frequently assigned cluster. *Scallop* has three key steps: 1) Bootstrapping, 2) Mapping between cluster labels across bootstrap iterations, and 3) Computation of the membership score.

#### Bootstrapping

*Scallop* runs a community-detection algorithm (default: Leiden (***Traag et al., 2019***)) on subsets of cells drawn from the original dataset. The subsets are selected randomly with replacement from the whole population (the seed can be defined by the user). The number of cells to be selected on each bootstrap iteration is computed through the fraction of cells user-defined parameter `frac_cells` (default: 0.95). The community detection algorithm is run `n_trials` times (default: 30). An additional clustering is run with all the cells (`frac_cells=1`) for it to be used as a reference in the mapping stage. A bootstrap matrix (`n_cells × n_trials`) is obtained that contains the cluster labels that have been assigned to each cell on each bootstrap iteration. The cluster labels are the ones obtained from the python implementation of Leiden through the *Scanpy* function `sc.tl.leiden()` and are numbered from ‘0’ to *n* according to the size of the cluster, i.e., the cluster with the highest number of cells is assigned the label ‘0’, the second most abundant is assigned the label ‘1’, and so on. Since the subset of cells used in each run is different, clustering results vary from run to run and labels are not comparable between bootstrap iterations.

#### Cluster relabeling

In order to compare between cluster assignments from different bootstrap iterations, cluster identities need to be relabeled. A contingency table is computed between each clustering solution in the bootstrap matrix and a reference clustering, which was obtained by running the community detection algorithm on all the cells. From the original bootstrap matrix, we obtain a relabeled bootstrap matrix. The assumption is made that if cluster A from bootstrap iteration *i* and cluster B from bootstrap iteration *j* have a large number of cells in common, then they should have the same label. In order to find the mapping between clusters, an overlap score matrix is computed for every column in the bootstrap matrix against the reference labels. The overlap score (*S*) between cluster A from the reference clustering solution (*A*^*ref*^) and cluster B from the *i*-th iteration (*B*^*i*^) is defined as follows:

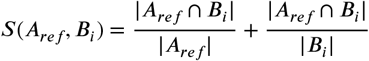

Where |*A*^*ref*^| and |*B*^*i*^| are the number of cells in the cluster *A* and *B* from the reference clustering solution and the *i*-th bootstrap iteration, respectively, and |*A*^*ref*^ ∩ *B*^*i*^| is the number of cells in common between the two clusters. The score is then [0-1]-scaled by dividing it by the maximum score: 2. The maximum score would correspond to a total overlap between the two clusters.

The score is computed for every pair of clusters between the reference solution and each of the bootstrap iterations to obtain a contingency matrix (*n*_*clusters*^*ref*^ × *n*_*clusters*^*i*^). In order to find the optimal mapping between the two clustering solutions, we search for the permutation of the columns that maximizes the trace of the contingency matrix. We do this by using Munkres, a Python implementation of the Hungarian method (***Munkres, 1957***).

As the reference clustering solution is computed on the whole dataset but each of the bootstrap iterations is run on a subset of cells (*frac*_*cells*), the number of clusters obtained in each iteration might not be equal to the number of clusters in the reference. In order to deal with this, we consider three cases:

1. The number of clusters in the reference clustering solution is equal to the number of clusters obtained in the *i*-th bootstrap iteration. This case is dealt with easily, as the Hungarian method yields a 1:1 mapping between the two clustering solutions.

2. Fewer clusters are obtained in the *i*-th bootstrap iteration than in the reference solution. This may happen if one or more clusters from the reference are merged into a single cluster in a bootstrap iteration. In this case, a 1:1 mapping is obtained, but one or more of the cluster labels from the reference clustering remain unused.

3. More clusters are obtained in the *i*-th bootstrap iteration than in the reference solution. This may happen if one cluster from the reference is further divided into two or more subclusters in a bootstrap iteration. A 1:1 mapping is obtained, but one or more clusters from the bootstrap iteration remain unmapped. Usually, this means than a cluster from the reference solution was divided into two or more subclusters in the bootstrap iteration. In this case, the subcluster with the largest overlap degree with one of the clusters in the reference clustering solution, receives its label. The other subcluster remains unmapped. When this happens, those clusters are flagged as *unmapped* until the end of the bootstrapping process. Then, an additional mapping step is carried out between the *unmapped* clusters from all bootstrap iterations. This is done by creating an overlap score matrix similar to the one created in the mapping process and searching for the permutation of the columns that maximizes its trace, using the Hungarian method. In order to avoid spurious mappings between unrelated *unmapped* clusters, a minimum overlap score of 0.1 is imposed for two *unmapped* clusters to be renamed as the same cluster.

#### Computation of the membership score

*Scallop* computes three different membership scores: a frequency score (‘freq’), an entropy score (‘entropy’) and a Kullback-Leibler divergence score (‘KL’). We use the frequency score here as it yields results that are consistent to the ones obtained with the other two alternative scores, and its meaning is more intuitive than those of the two alternative methods. The frequency score is computed as the fraction of bootstrap iterations where a cell was assigned to the most frequently assigned cluster label. In order for the score to take values between 0 and 1, only the cells selected in each bootstrap run are considered as the total number of cells. More information on the calculation of the entropy and the Kullback-Leibler scores can be found in the *Scallop* documentation.

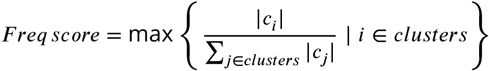

### Validation of *Scallop*

In order to validate our method for transcriptional noise quantification, we conducted three analyses:

1) we graphically evaluated the output of *Scallop* on a dataset of human T cells, 2) we analyzed its robustness to input parameters, and 3) we studied the relationship between membership and robust marker expression, using a PBMC dataset.

#### Stable and unstable cells in the 8K human T cells

We downloaded a 23,766 PBMC dataset from from 10X Genomics. We ran the standard processing pipeline including highly variable gene detection, dimensionality reduction through PCA and UMAP, and clustering. We annotated the dataset according to PBMC marker expression and selected the cluster of T lymphocytes. We obtained a dataset of 8,278 cells. We ran the processing pipeline on the T lymphocyte dataset and obtained three main clusters of cells, which we annotated as *0/CD4+ T cells, 1/CD4+ T cells* and *2/CD8+ T cells* according to their expression of the gene markers *CD3C, CD3D, CD3E, CD4, CD8A* and *CD8B*. Then, we calculated the whole transcriptome-based Euclidean distance to the cell subtype mean (*euc_dist)*), the invariant gene-based Euclidean distance to the T cell mean (*euc_dist_tissue_invar*), the *Scallop* noise as 1-membership (*scallop_noise*) and the GCL per T subtype. We selected the 10% most stable and 10% most unstable cells as those having the lowest and highest noise scores according to two methods: *euc_dist* and *scallop_noise*.

#### Robustness to input parameters

We selected a set of five scRNAseq datasets of various sizes and depths (Table 1). Three datasets were taken from published scRNAseq studies (***Paul et al., 2015; Moignard et al., 2015; Joost et al., 2016***) and two were from 10X Genomics (PBMC3K, Heart10K). We computed the membership scores of all the cells in five datasets 100 times, on a range of bootstrap iterations (`n_trials`), fraction of cells used in each iteration bootstrap (`frac_cells`) and resolution (res) values. We then computed the median correlation distance between the 100 runs of *Scallop* with each set of parameters. We used the spatial.distance.correlation method from Scipy to compute the correlation distance.

#### Statistical significance of differential expression of PBMC markers

The assessment of the cell-to-cell variability associated with aging using *Scallop* relies on the assumption that cell-to-cell variability is caused by transcriptional noise, and that it can be measured by evaluating cluster stability. We checked our assumption by comparing the transcriptomic profiles of the cells that had a high and a low membership score (measured using *Scallop*). *Stable* cells should have a more robust expression of cell type markers than the *unstable* cells. We downloaded the PBMC 3K datataset from 10x Genomics. After running the standard processing pipeline, we ran *Scallop* on the dataset and selected the most stable and most unstable half of the cells within each annotated cluster. For each cell type, we defined the most stable cells as those with a membership score greater than the median membership score of that cell type. Hence, we compared two sets of cells (stable vs unstable) of the same size and we analyzed the effect size and statistical significance of a routine downstream analysis (differential expression) when given each of the sets as input. We computed the 100 most differentially expressed genes (DEGs) between each cell type and the rest of the cells using 1) all cells, 2) only the stable cells and 3) only the unstable cells. B cells and megakaryoctes were excluded from the analysis as the former were highly stable (so we could not compare between the stable and the unstable fraction) and the latter consisted of very few cells. We compared the distribution of log-fold changes and *p*-values associated with those DEGs when using only the stable, only the unstable and all the cells.

### Single-cell RNA sequencing data processing

#### 2,5K human aging pancreatic cells

The raw count matrices and the metadata files from ***Enge et al***. were downloaded from the Gene Expression Omnibus (accession number: GSE81547). The separate GSM files were merged into a single raw count matrix and processed them using the following pipeline in Scanpy (***Wolf et al., 2018***): filtering of low quality cells and genes, normalization, log-transformation of counts, PCA, batch-effect correction using harmony (***Korsunsky et al., 2019***), Leiden community detection (resolution=1.0) and UMAP dimensionality reduction. 11 clusters were obtained and annotated using the expression of the markers *INS* (*β* cells), *GCG* (*α* cells), *SST* (*δ* cells), *PRSS1* (acinar cells), *PROM1* (ductal cells), *PPY* (PP cells) and *THY* (mesenchymal cells). Donors were classified into three categories as in the original work by ***Enge et al***..: “pediatric” (0-6 years old), “young” (21-22 years old) and “old” (38-54 years old). Samples from pediatric donors were not used in the aging analysis (see Inclusion Criteria 3).

#### 1,5K murine aging CD4+ T cells

We downloaded the raw data and metadata files from (***Martinez-Jimenez et al., 2019***) from the authors’ GitHub. We created an annData object with the raw count matrix and the metadata (mouse strain, age-group, stimulus, individual and cell type). We identified and flagged the counts corresponding to ERCC spike-in controls. We ran a standard processing pipeline: filtering out low quality cells and genes, normalization and log-transformation of counts, selection of highly variable genes, batch-effect correction between mouse strains (*Mus musculus domesticus* and *Mus musculus castaneus*) and dimensionality reduction was conducted (PCA and UMAP).

#### 14,8K murine aging lung cells

We downloaded the raw count matrix and the metadata file from (***Angelidis et al., 2019***) from the Gene Expression Omnibus (accession number: GSE124872). We created an annData object with the raw count matrix and the available metadata (cell type annotation, age group, cluster and mouse). We ran a standard processing pipeline: quality control, normalization and log-transformation of counts, selection of highly variable genes, batch-effect correction between individual mice using bbknn (***Park et al., 2018***) and dimensionality reduction (PCA and UMAP). In our analysis, we used the cell type annotations provided by the authors. We also annotated the rest of the murine aging lung datasets using their annotation as a reference. In order to do that, we computed the DEGs between each lung cell type and the rest of the dataset to obtain a set of gene markers for each cell type. We then used those markers to annotate the rest of the datasets using scoreCT (***Seninge, 2020***).

#### 90,6K murine aging lung, spleen and kidney cells

We downloaded the raw count matrices and the metadata files from ***Kimmel et al***. from the Gene Expression Omnibus (accession number: GSE132901). We selected lung samples (30,255 cells) and excluded kidney and spleen samples. We discarded the two samples from the individual Y1, as they showed a very different count distribution to the rest of the samples (see Appendix 1). We created an annData object with the count matrix and the metadata (sample, tissue, age, mouse). We ran a standard processing pipeline: quality control, normalization and log-transformation of counts, highly variable gene selection, batch-effect correction between individual mice using bbknn (***Park et al., 2018***), dimensionality reduction (PCA and UMAP) and Leiden clustering (***Traag et al., 2019***) with high resolution value (resolution = 4), so that we obtained a very granular clustering solution. We obtained 52 clusters and we annotated them by projecting the cell type identity labels from the ***Angelidis et al***. dataset, using the automated cell type annotation tool scoreCT (***Seninge, 2020***). We checked that cells clustered primarily according to their cell type, meaning no important batch effects were present in the final datasets, and that clusters expressed the cell type markers expected according to their assigned cell type labels (see Appendix 1).

#### 731 murine aging dermal fibroblasts

The count matrix and metadata from ***Salzer et al***. were downloaded from the Gene Expression Omnibus (accession number: GSE111136). A standard processing pipeline (quality control, normalization, log-transformation, HVG detection, PCA, neighbor computation and UMAP dimensionality reduction) was applied to the dataset.

#### 22,1K human aging skin cells

We downloaded raw count matrices from ***Solé-Boldo et al***. from the Gene Expression Omnibus (accession number: GSE130973). We ran a standard preprocessing pipeline on the count matrix: quality control, normalization, log-transformation, HVG detection, PCA, neighbor computation and UMAP dimensionality reduction. We used the original cell type labels provided by the authors.

#### Tabula Muris Senis lung datasets

The 3,2K TMS FACS-sorted and the 4,4 TMS droplet lung cell datasets were downloaded from figshare. A standard preprocessing pipeline was run on the two datasets, and cluster labels were harmonized with the rest of the murine aging lung datasets by using the genes differentially expressed between cell types from the Angelidis dataset as input for the automated cell-type annotation through scoreCT (***Seninge, 2020***)

#### Human Lung Cell Atlas

We downloaded the full lung and blood 10X dataset from the HLCA (***Travaglini et al., 2020***) from Synapse (ID: syn21041850). The original dataset consists of lung samples from 3 patients: a 46 years old male donor (donor 1), a 51 years old female donor (donor 2) and a 75 years old male donor (donor 3). The composition of the samples was not equivalent across donors: there were two samples from donor 1 (distal and medial), three samples from donor 2 (blood, distal and proximal) and two samples from donor 3 (blood and distal). Thus, we selected the distal sample from the three donors and obtained a dataset of 18,542 cells from donor 1, 16,903 cells from donor 2 and 7,524 cells from donor 3. We subsampled 7,524 cells from each of the donors in order to correct for the age-group imbalance, and obtained a dataset of 22,572 lung cells. We used this balanced dataset of distal samples from the three donors to create two datasets. On the one hand, we selected all lung cells from donors 1 and 3 (46 years old and 75 years old, all male) in order to create the 15,048 aging lung cell dataset used in the noise and enrichment analysis. On the other hand, we selected all alveolar macrophages from the three donors in order to create the 11,484 alveolar macrophage dataset.

#### Human Aging Lung

We downloaded the mammalian aging lung dataset by ***Raredon et al***. from the Gene Expression Omnibus (accession number: GSE133747). The original dataset consists of human, pig, mouse and rat samples. We selected human samples and ran the preprocessing and quality control pipeline on them: normalization, log-transformation, selection of highly variable genes, batch-effect correction between donors using harmony (***Korsunsky et al., 2019***), computation of the nearest neighbor graph and Leiden clustering (***Traag et al., 2019***). The resulting dataset consisted of 17,867 cells from human male and female donors aged 21 to 88 years. We then projected the cell type labels from the human lung atlas onto the Raredon dataset by computing the DEGs between cell types in the Human lung atlas dataset and using the first 300 DEGs to identify equivalent cell types in the Raredon dataset and projecting those onto the Raredon dataset using the unsupervised cell type annotation tool scoreCT (***Seninge, 2020***). We identified 24 lung cell types from the HLCA. After using the cells from the 14 the human donors in the annotation step, we selected a set of 9 donors in order to obtain a balanced aging dataset, using the following inclusion criteria: 1) Donors contributing with very few cells were excluded (GSM4050113 and GSM4050107 consisted of 116 and 211 cells, respectively), 2) middle-aged donors were discarded in order to better explore the effects of aging,

3) donors were selected to ensure sex-stratification, 4) we sought to obtain a balanced dataset in terms of age-group sizes. The final dataset consisted of 15,852 lung cells from 9 female and male human donors. We defined the age categories as young (21, 22, 32, 35 and 41 years old) and old (64, 65, 76 and 88 years old). The composition of the dataset was 7,263 young (46%) and 8,589 old cells (54%).

### Age-related change in transcriptional noise

#### Measuring age-related loss of cell type membership

To facilitate comparison with regard to cell type annotation, we harmonized the labels so that the four datasets were annotated using the cell identities originally defined by ***Angelidis et al***.. Then, we measured transcriptional noise as 1 − membership to cell type clusters in the young and old fractions of each dataset. We then measured the age-related difference in transcriptional noise per cell type by calculating the differences in median noise between the old and the young fraction for each lung cell type. In order to compare between the young and the old fraction of cells, each dataset was split into two datasets according to the age groups (“young” and “old”), and the highly variable gene detection and dimensionality reduction (PCA, batch-corrected neighbor detection using harmony, and UMAP) steps where run again on each set of cells. Then, *Scallop* was run on each set of cells separately, using Leiden as the community detection method and using the following parameter values: `frac_cells=0.8, n_trials=30`. This was done on a range of resolution (`res`) values between 0.1 and 1.5, with a step of 0.1, and the membership scores obtained for each cell were averaged over all these resolution values in order to smooth the effect of clustering granularity on the membership scores. We used the *freq* membership score, defined as the frequency of assignment of the most frequently assigned cluster label per cell.

### Age-related cell type enrichment

Changes in cell type abundance associated to aging were evaluated using binomial GLMs ***McCullagh and Nelder*** (***1989***). For each dataset, a binomial GLM was fitted to estimate the proportion of each cell type across all samples by treating each individual mouse as a replicate. First, the relative abundance of each cell type (*N_ct*) and the relative abundance of the rest of the cell types taken together (*N_other*) was computed. Then, a binomial GLM was fitted to these pairs of observations (*N_ct, N_other*) to estimate the proportions of cell types across samples by accounting for variation associated to sample origin (mouse) and to sample age (young vs old), and estimated marginal means (***Searle et al., 1980***) were computed using the R package *emmeans*. Odds ratios between “Young” and “Old” samples were computed for each cell identity.

## Data availability

The Angelidis dataset was downloaded from the Gene Expression Omnibus (GSE124872). The original Kimmel dataset (consisting of kidney, spleen and lung samples) was downloaded from the Gene Expression Omnibus (GSE132901). The preprocessed and annotated files for the TMS datasets were downloaded from figshare (***Pisco, 2020***). The final version of the datasets as well as the reproducible Jupyter notebooks with all the analyses used in this study can be found in figshare (10.6084/m9.figshare.14981181).

## Code availability

The *Decibel* and *Scallop* repositories can be found at https://gitlab.com/olgaibanez/decibel and https://gitlab.com/olgaibanez/scallop, respectively. The reproducible Jupyter notebooks with the analyses carried out in this study can be found in figshare (10.6084/m9.figshare.14981181).

## Funding

This work was supported by grants from Instituto de Salud Carlos III (AC17/00012 and PI19/01621), cofunded by the European Union (European Regional Development Fund/European Science Foundation, Investing in your future) and the 4D-HEALING project (ERA-Net program EracoSysMed, JTC-2 2017); Diputación Foral de Gipuzkoa; Ministry of Science and Innovation of Spain; and PID2020-119715GB-I00 funded by MCIN/AEI/10.13039/501100011033 and by “ERDF A way of making Europe”. OI-S received the support of a fellowship from “la Caixa” Foundation (ID 100010434; code LCF/BQ/IN18/11660065), and from the European Union’s Horizon 2020 research and innovation programme under the Marie Skłodowska-Curie grant agreement No. 713673. AMA was supported by a Basque Government Postgraduate Diploma fellowship (PRE_2020_2_0081).

## Acknowledgments

We thank Iñaki Inza for his thorough revision of the manuscript, Laura Yndriago for her feedback, and Sandra Fuertes for useful discussions. We thank Valentine Svensson for support with the application of GLMs to cell type abundance analysis of scRNAseq data.

## Author contributions

Conceptualization: OI-S; Funding Acquisition: MJA-B, OI-S, AMA; Investigation: OI-S, AMA, AI; Methodology: OI-S, AMA; Project Administration: AI, MJA-B; Resources: MJA-B; Software: OI-S, AMA; Supervision: AI; Visualization: OI-S, AMA; Writing - Original Draft Preparation: OI-S, AI; Writing - Review and Editing: OI-S, AMA, MJA-B, AI.

## Appendix 1

**Appendix 1 Table 1.** Datasets used in the technical validation of *Scallop*. Number of cells, number of genes, median number of genes per cell and number of estimated cell populations in each dataset.

| Dataset | n_cells | n_genes | median_genes_per_cell | n_populations |
| --- | --- | --- | --- | --- |
| PBMC3K | 2700 | 32738 | 817 | 12 |
| Joost et al. | 1422 | 6410 | 1941 | 17 |
| Paul et al. | 2730 | 3451 | 872 | 14 |
| Moignard et al. | 3934 | 42 | 42 | 9 |
| Heart10K | 7713 | 11765 | 2035 | 26 |

## Appendix 2

**Appendix 2 Table 1.** Seven scRNAseq studies of different tissues where age-related increase in transcriptional noise was measured. The number of cells (*N. cells*) in the table is the size of the dataset prior to quality control. The *Noise* column states whether an increase in transcriptional noise was reported in some/all cell types in the original articles. The *Scope* column summarizes the cell types where age-related increase in transcriptional noise was reported. The *Method* column specifies how transcriptional noise was measured in the original articles.

| Dataset | Tissue | Organism | N. cells | Noise | Scope | Method |
| --- | --- | --- | --- | --- | --- | --- |
| *Enge et al.* | Pancreas | Human | 2,544 | Yes | In Beta cells. | 1) Biological over technical variation, 2) whole transcriptome-based Euclidean distance to cell type mean, 3) invariant gene-based Euclidean distance to cell type mean. |
| *Martinez-Jimenez et al.* | CD4+ T cells | Mouse | 1,513 | Yes | Single cell type studied. | Percentage of cells expressing the core activation program. |
| *Angelidis et al.* | Lung | Mouse | 14,813 | Yes | In most cell types. | Distance to cell type mean. |
| *Kimmel et al.* | Lung, spleen, kidney | Mouse | 30,255<br>30,512<br>29,815 | Yes | In many cell types. | 1) Overdispersion of genes, 2) invariant gene-based Euclidean distance to cell type mean, 3) whole transcriptome-based Manhattan distance to cell type mean. |
| *Ximerakis et al.* | Brain | Mouse | 37,069 | No | Differences in magnitude and directionality between cell types. | Coefficient of variation of 1) all genes, 2) mitochondrial genes, 3) ribosomal genes. |
| *Salzer et al.* | Dermal fibroblasts | Mouse | 731 | Yes | Single cell type studied. | Compactness of clusters on PCA plot. |
| *Solé-Boldo et al.* | Skin | Human | 22,142 | Yes | In dermal fibroblasts. | Less clear GO annotations. |

## Appendix 3

**Appendix 3 Table 1.** The general criteria for inclusion in the aging datasets used in this study was to include all samples from young and old individuals and to exclude newborn or pediatric individuals, as we did for the human pancreatic cell dataset (***Enge et al., 2017***) and the murine dermal fibroblast dataset (***Salzer et al., 2018***). Care was taking to make all aging datasets sex-balanced. This was not possible for some datasets, as they consisted of same-sex individuals. However, same-sex datasets were included in our study as sex could not be a confounding factor in the aging analysis.

| Dataset | Inclusion criteria | Number of individuals | Number of cells | Ages |
| --- | --- | --- | --- | --- |
| *Enge et al.* | All samples except those from pediatric individuals (0-6 y old) | Young: 2<br>Old: 3 | Young: 791<br>Old: 1,023 | Young: 21 and 22 y<br>Old: 38, 44 and 58 y |
| *Martinez-Jimenez et al.* | Whole dataset. | Young: 9<br>Old: 12 | Young: 532<br>Old: 981 | Young: 3 m<br>Old: 24 mo |
| *Angelidis et al.* | Whole dataset. | Young: 8<br>Old: 7 | Young: 7,644<br>Old: 6,526 | Young: 3 m<br>Old: 24 m |
| *Kimmel et al.* | Lung samples from all mice except Y1. | Young: 3<br>Old: 3 | Young: 13,352<br>Old: 12,998 | Young: 7 m<br>Old: 24 m |
| *Ximerakis et al.* | Whole dataset. | Young: 8<br>Old: 8 | Young: 16,028<br>Old: 21,041 | Young: 2-3 m<br>Old: 21-23 m |
| *Salzer et al.* | All samples except those from newborn mice. | Young: 4<br>Old: 4 | Young: 329<br>Old: 332 | Young: 2 m<br>Old: 18 m |
| *Solé-Boldo et al.* | Whole dataset. | Young: 2<br>Old: 3 | Young: 8,316<br>Old: 13,826 | Young: 25 and 27 y<br>Old: 53, 69 and 70 y |

## Appendix 4

**Appendix 4 Figure 1.**
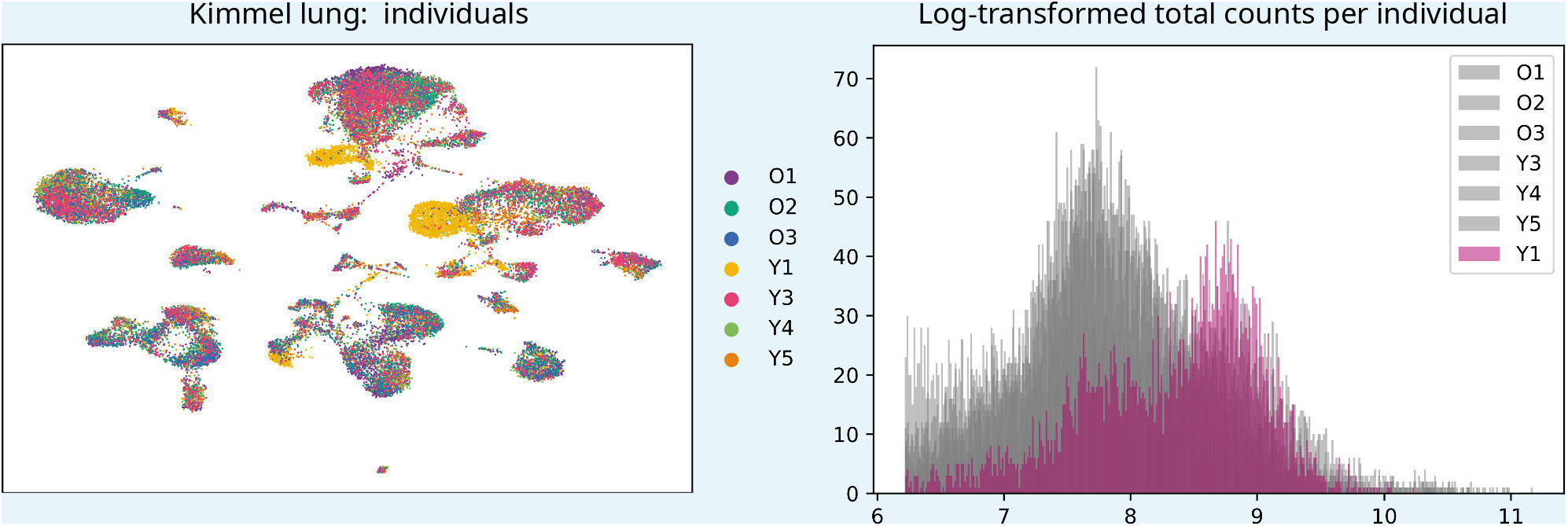
UMAP plot showing the samples from the seven individuals present in the Kimmel lung dataset. Even though most cells cluster together according to their cell type rather than by individual, samples from donor Y1 cluster together. We observed that there was a big batch effect between this and the rest of the individuals. **Histogram showing the log-transformed total number of counts/cell per individual mice**. The distribution of counts/cell of the samples from mouse Y1 is very different to the rest of the samples. This difference could not be overcome using the batch-effect correction tool *bbknn*. Downsampling the counts so that the number of counts/cell was balanced across individual mice did not solve the problem either. Therefore, we decided to discard the samples Y1L1 and Y1L2.

## Appendix 5

**Appendix 5 Figure 1.**
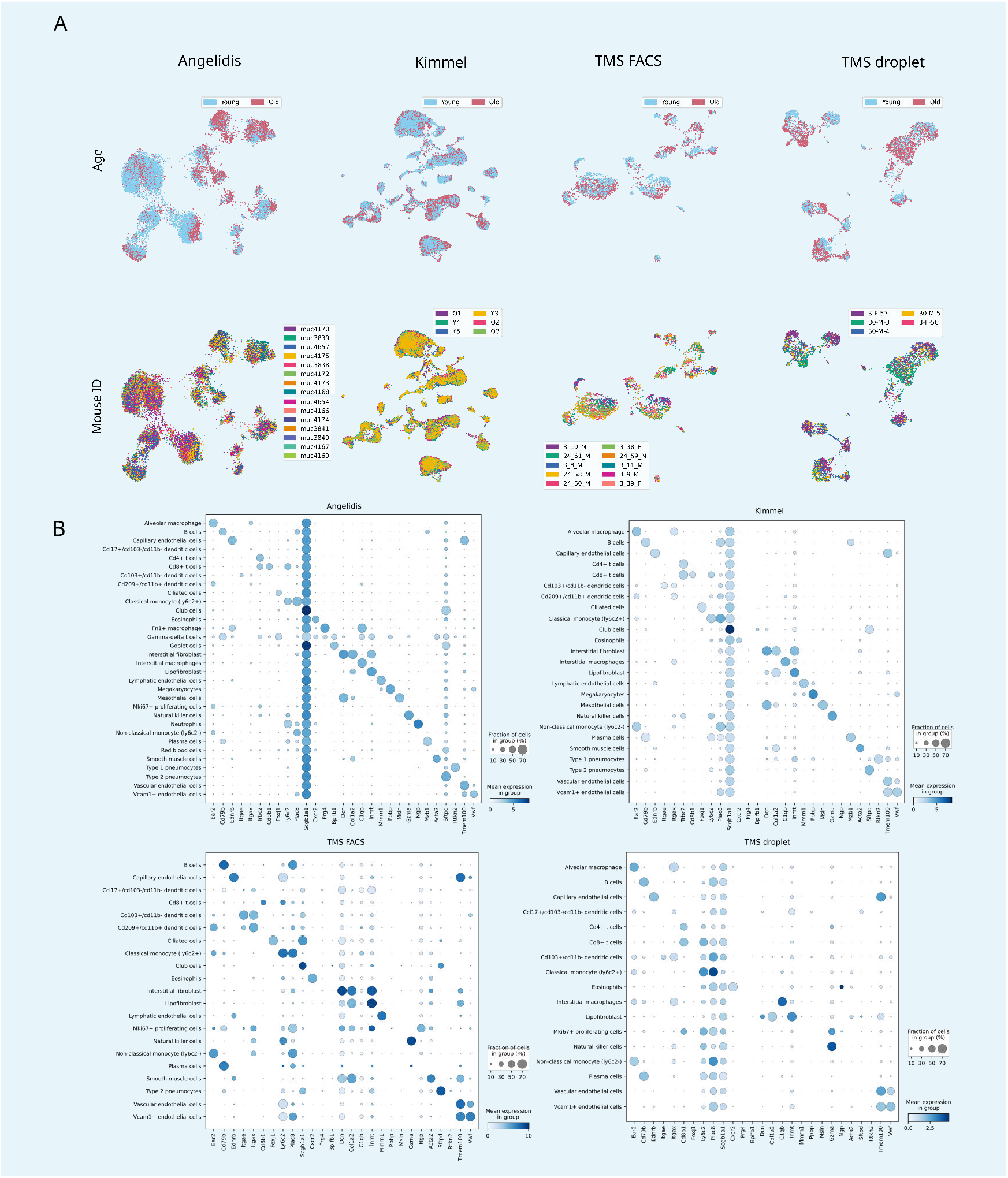
(A) There are no mouse- or age-related batch effects. UMAP plots of the four aging lung datasets showing the age and mouse labels. Cells cluster according to their cell type rather than to their age group or individual mouse. **(B) Expression of lung cell type markers by each annotated cluster**. The dotplots show the expression of the cell type markers from ***Angelidis et al***. on the four annotated lung datasets. The size of the dots represents the fraction of cells expressing one particular marker in the group of cells assigned a particular cell type label. The color represents the level of expression of the marker in that group averaged over the cells that have a positive expression of that marker.

## Appendix 6

**Appendix 6 Table 1.** Number of cells, sex and age composition of the human aging lung datasets.

| Dataset | N. young cells | N. old cells | N. young donors | N. old donors | Total cells |
| --- | --- | --- | --- | --- | --- |
| *Raredon et al.* | 7,263 | 8,589 | 3 females: 21, 32, 41 years old<br>2 males: 22, 35 years old | 2 females: 76, 88 years old<br>2 males: 64, 65 years old | 15,852 |
| *Travaglini et al.*<br>(HLCA) | 7,524 | 7,524 | 1 male: 46 years old | 1 male: 75 years old | 15,048 |

**Figure 1–Figure supplement 1.**
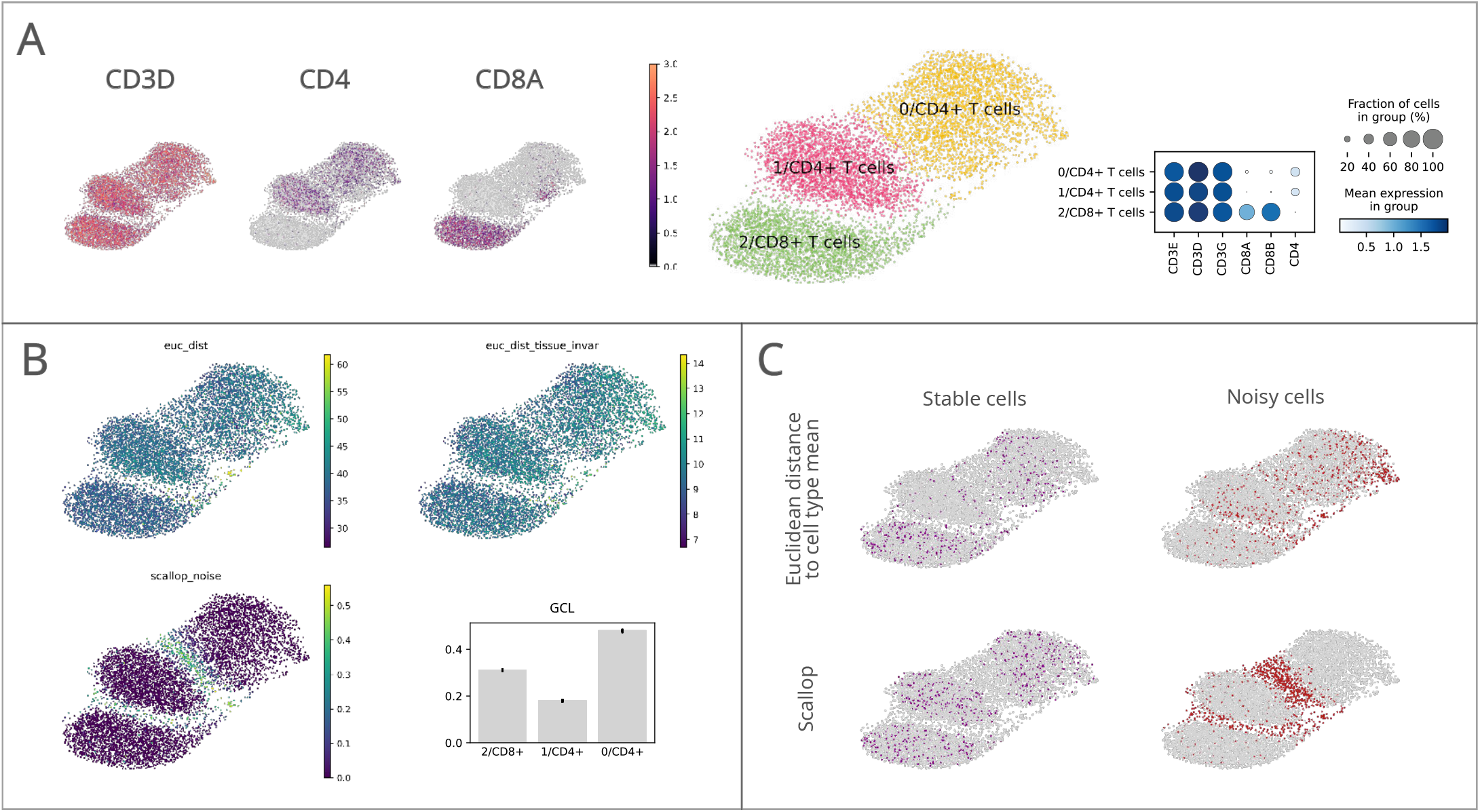
Performance of *Scallop* in comparison to preexisting methods for the quantification of transcriptional noise. The different methods were tested on a dataset of 8,278 human T lymphocytes. (A) UMAPs and dotplot showing *CD3, CD4* and *CD8* marker gene expression per cluster. (B) Representation of transcriptional noise levels, as measured by using two distance-to-centroid methods (*euc_dist* and *euc_dist_tissue_invar*), 1 - membership (*scallop_noise*) and Global Coordination Level (*GCL*). (C) The 10% most stable (purple) and 10% most unstable (red) cells are represented on the UMAP plots for *Euclidean distance to cell type mean* (top row) and *Scallop* methods (bottom row), respectively.

**Figure 1–Figure supplement 2.**
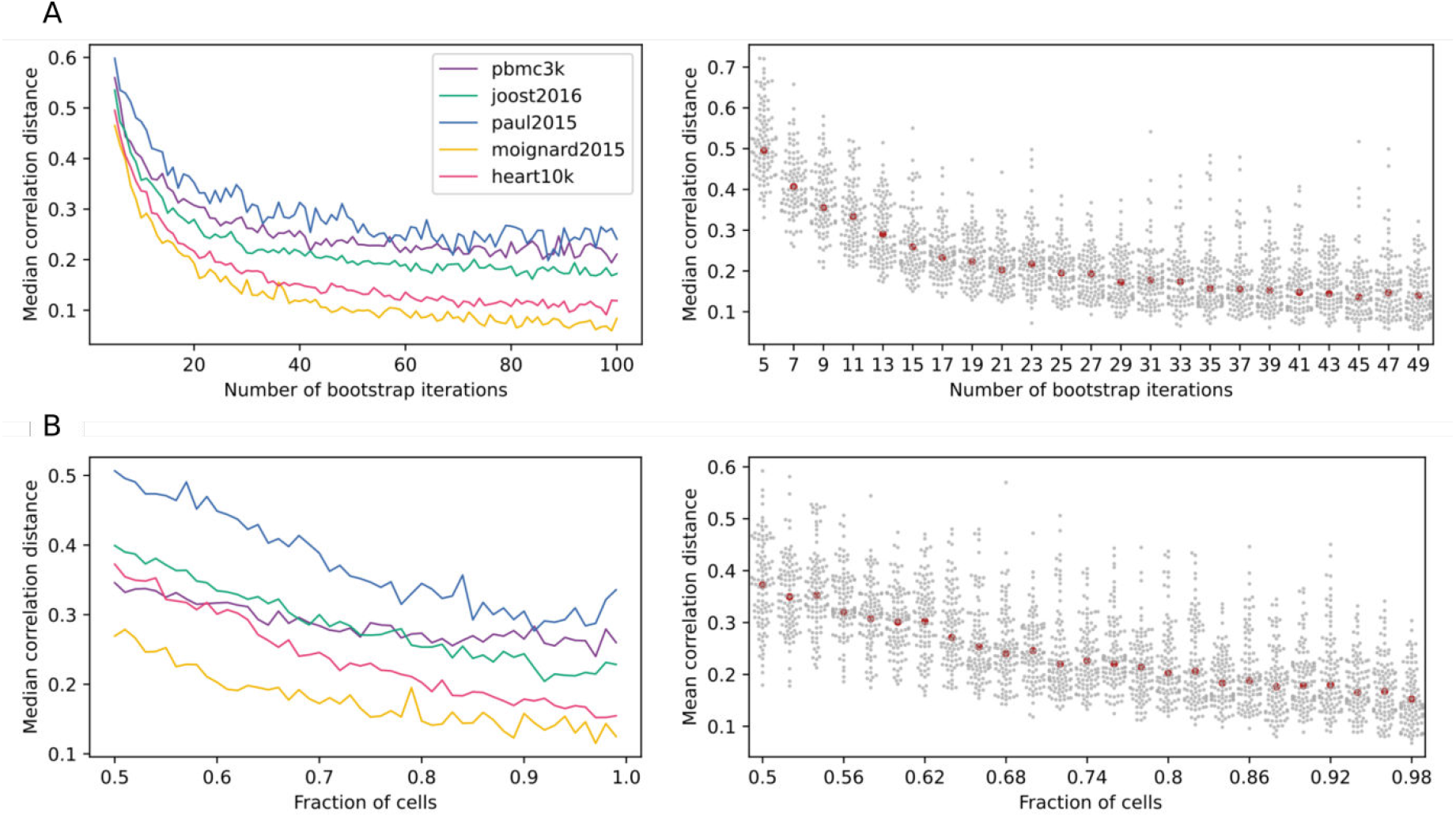
*Scallop* robustness in relation to input parameters. The plots on the left show the median correlation distance between membership scores of different runs of *Scallop* against (A) the number of trials, (B) the fraction of cells used in each bootstrap and (C) the resolution given to the clustering method (Leiden) in five independent scRNAseq datasets (PBMC3K, ***Joost et al. (2016***); ***Paul et al. (2015***); ***Moignard et al. (2015***), Heart10K). The median correlation distance was computed over 100 runs of *Scallop*. The swarmplots on the right show the distribution of the correlation distances between membership scores against each of the input parameters for the heart10k dataset. The median is shown as a red point. While, for the sake of clarity, a random sample of 100 correlation distances is shown for each value of the parameter under study, the median was computed using all the correlation distances. *Scallop* membership scores converge as we increase the number of bootstrap iterations and the fraction of cells used in the clustering.

**Figure 1–Figure supplement 3.**
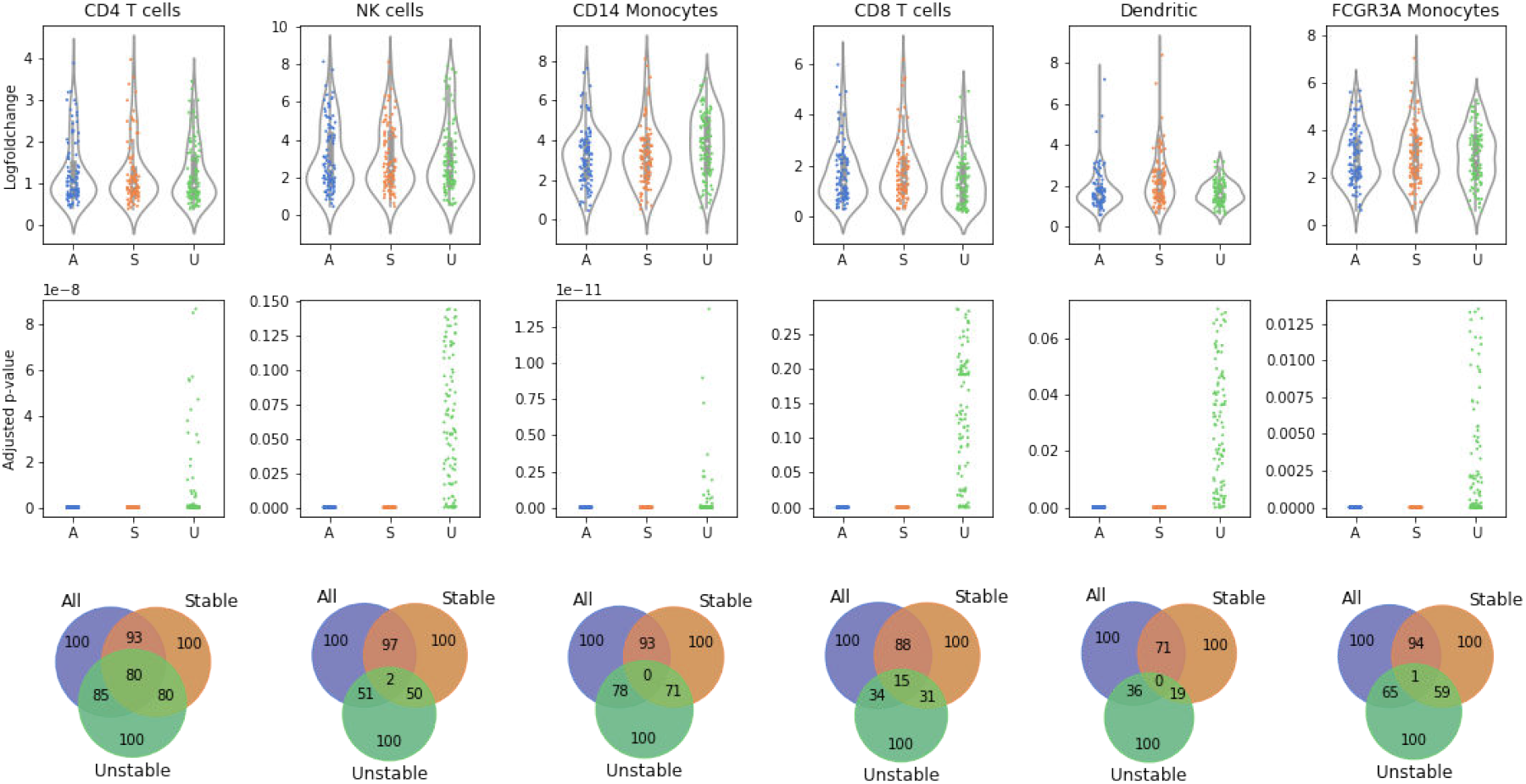
Stable cells as identified with *Scallop* are more representative of the cell type than unstable cells. Distribution of log-fold changes (top row) and adjusted *p*- values (middle row) of the first 100 differentially expressed genes (DEGs) between each cell type or subtype and the rest of the cells in six cell types and subtypes from the 10X PBMC3K dataset. The overlap between the DEGs found when using all of the cells, only the stable cells and only the unstable cells is also shown (bottom row). The adjusted *p*-values obtained with all the cells are equivalent to those obtained using only the most stable half of the cells. In contrast, the differential expression of many genes is not statistically significant when using the unstable half from each population. The overlap between the top 100 DEGs obtained is very high between the stable cells and all cells subsets, whereas DEGs obtained in unstable cells have a very low intersection with all cells.

**Figure 2–Figure supplement 1.**
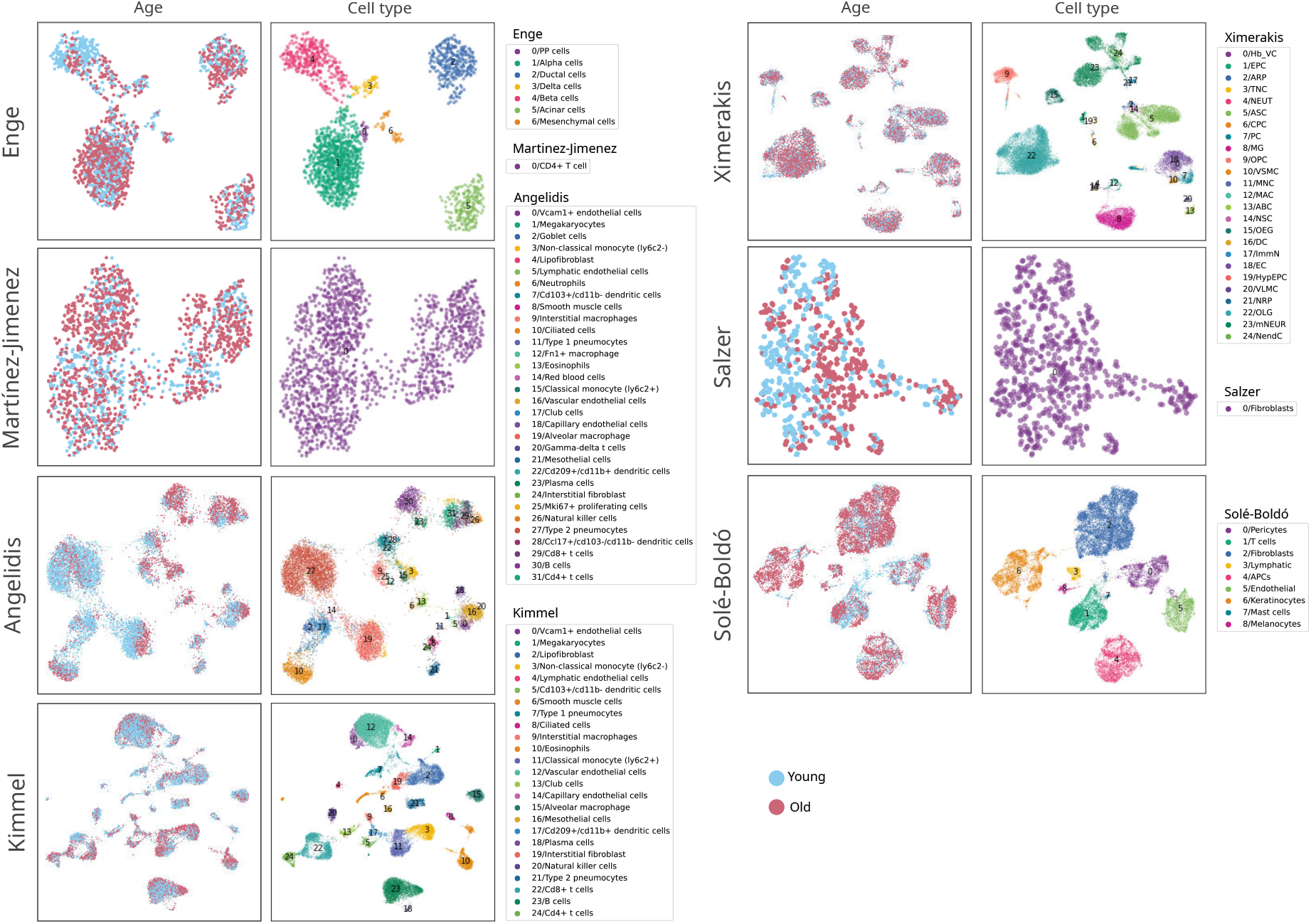
Composition of the seven scRNAseq datasets of aging used in this figure. UMAPs showing the age and cell type composition of the seven datasets used in the analysis of the age-related transcriptional noise at the tissue level. The UMAPs show the final composition of the datasets used in the experiment. The cell type annotations were obtained from the original authors in all datasets except Kimmel lung, where the labels from Angelidis were projected onto the dataset.

**Figure 2–Figure supplement 2.**
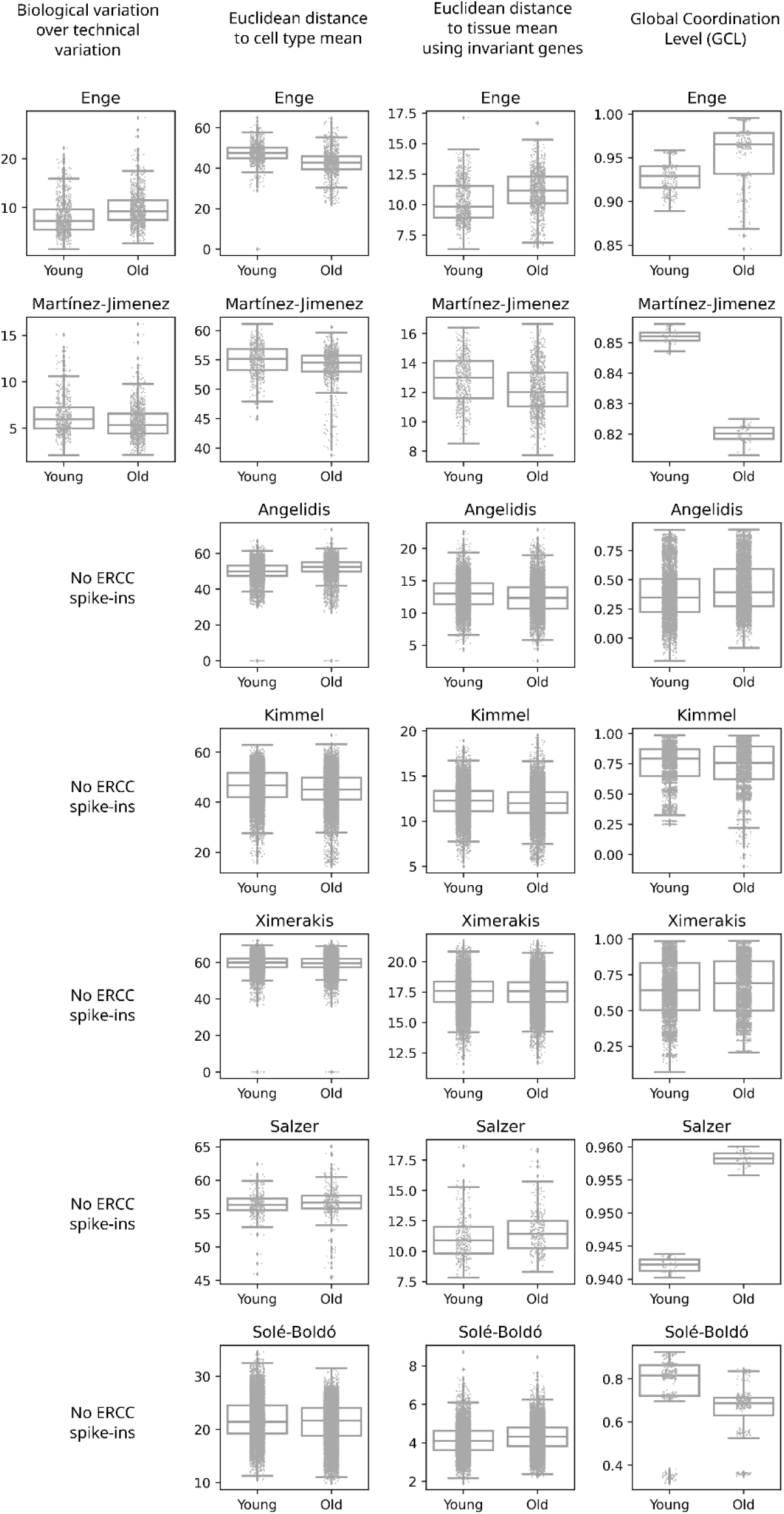
Measurements of transcriptional noise on seven scRNAseq datasets of aging using computational methods implemented in *Decibel*. Stripplots showing the distribution of noise values, as measured by the four alternative methods (Biological variation over technical variation, Euclidean distance to cell type mean, Euclidean distance to tissue mean using invariant genes, and Global Coordination Level - GCL) in the seven datasets used in the analysis of the age-related transcriptional noise at the tissue level. Boxplots and their whiskers represent the interquartile range (IR) and 1.5*IR respectively.

**Figure 3–Figure supplement 1.**
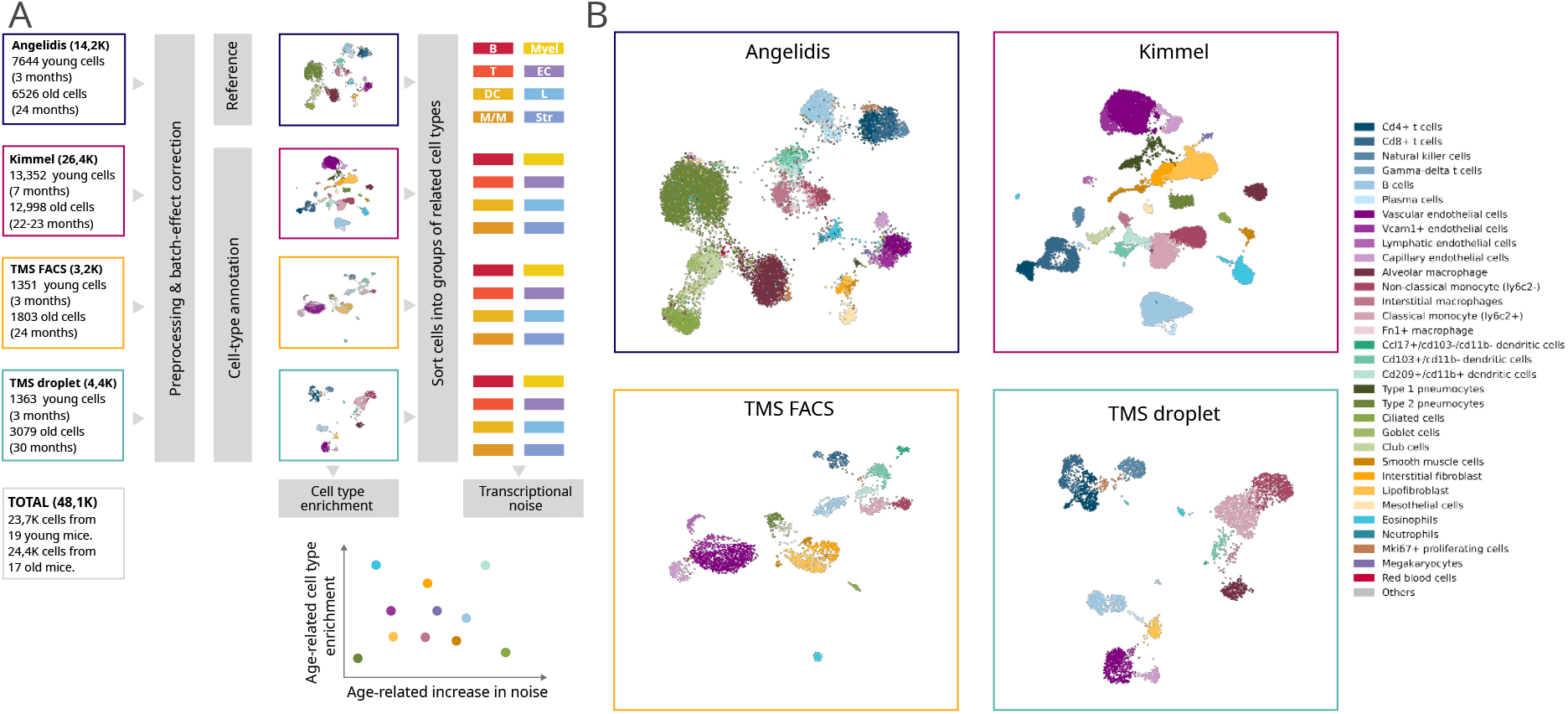
Composition of the four scRNAseq datasets of the murine aging lung used in this figure. (a) Experimental approach. Four murine aging lung datasets were preprocessed and cell type-annotated. The cell-type labels from Angelidis were used as a reference to annotate the rest of the datasets. Differences in cell-type abundance between young and old mice were quantified using GLMs. From each dataset, eight subsets of related cell-types were created to classify the 31 cell types into 8 categories, which were used as input for *Scallop* to analyze the differences in cell-to-cell variability. (b) Cell type-annotated mouse lung datasets. UMAP plots showing the four datasets with their cell type annotations.

**Figure 3–Figure supplement 2.**
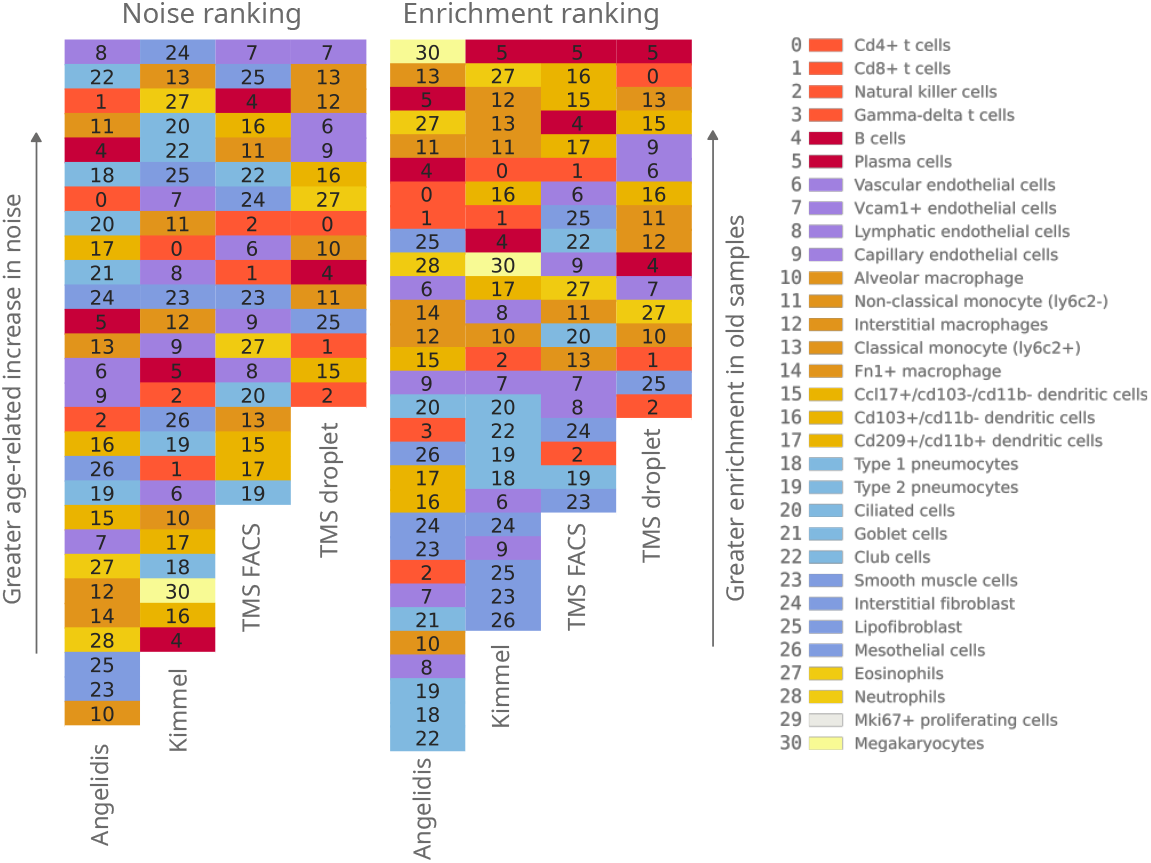
Qualitative ranking of murine aging lung cell types according to transcriptional noise and cell type enrichment. The 31 detected lung cell types were classified in the *Noise* ranking (left) according to their greater age-related increase in noise. They were also classified in the *Enrichment* ranking (right) according to their greater enrichment in old samples. Cell categories that were represented by fewer than 100 cells were excluded from the transcriptional noise evaluation, and therefore do not appear in the plot. Specific cell types are shown in the same color and with the same numbers as specified in the legend.

**Figure 3–Figure supplement 3.**
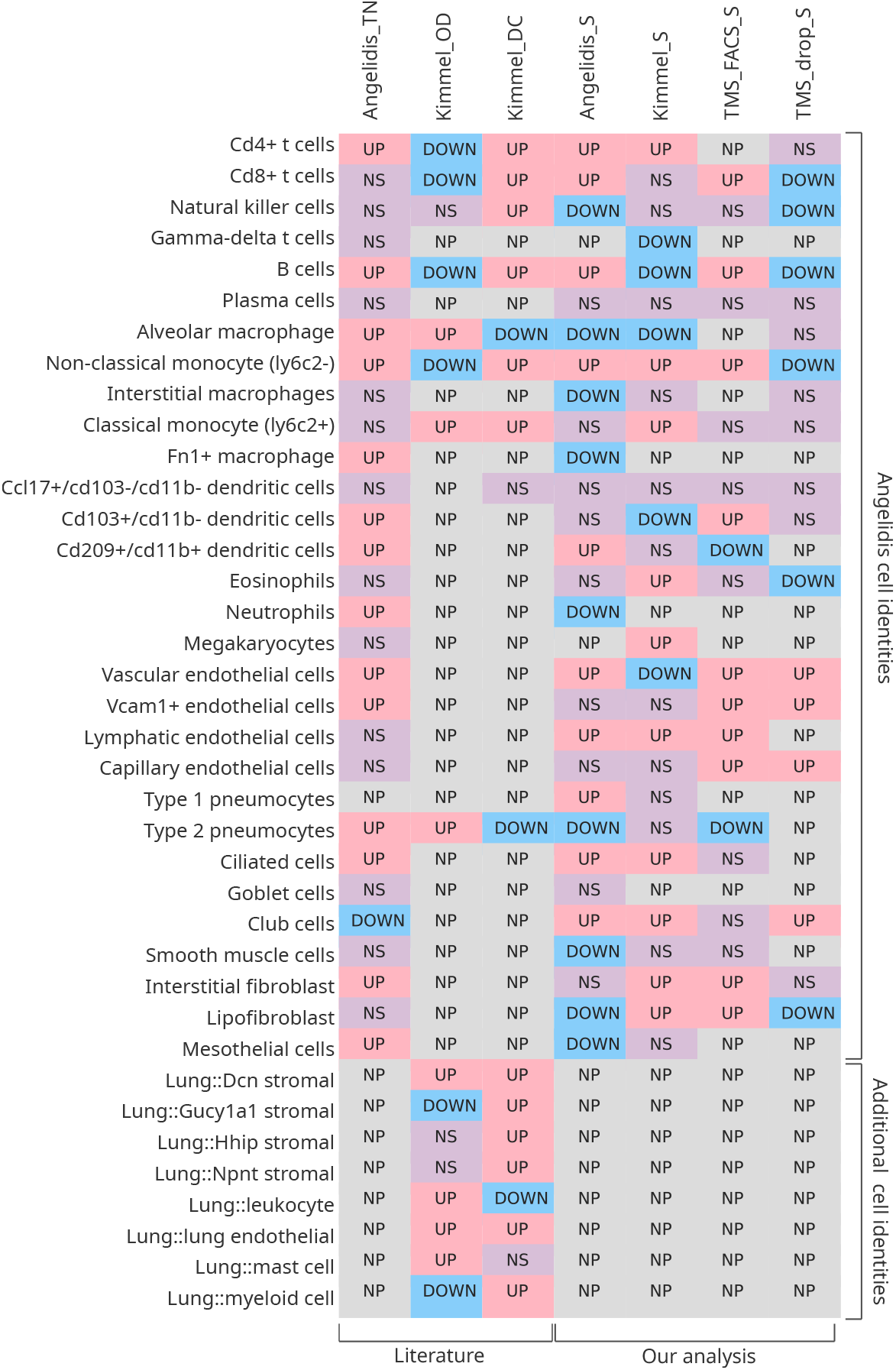
Comparison of the originally reported cell type-associated increase in transcriptional noise with the results obtained with *Scallop*. The content of the first three columns was drawn from the original publications (***Angelidis et al***.; ***Kimmel et al***.). More specifically, *Angelidis_TN* is the transcriptional noise per cell identity on the Angelidis dataset (from their figure 2); *Kimmel_OD* is the gene overdispersion per cell type on the Kimmel dataset (from their figure 2B); and *Kimmel_DC* is the cell-cell heterogeneity per cell identity measured as the Euclidean distance to the centroid of the cell identity for a particular age. Columns 4-7 summarize the results of our analysis of age-related loss of cell type identity in the murine lung. Specifically, *Angelidis_S, Kimmel_S, TMS_FACS_S and TMS_drop_S* report the transcriptional noise per cell identity on the four datasets, measured as the difference in median membership score between young and old individuals. The cell identities used are those drawn from Angelidis. Since some cell identities from Kimmel dataset did not have a 1:1 correspondence to the Angelidis cell identities, they are shown using their original notation at the bottom of the table (“Additional cell identities”). UP/DOWN: age- related increase/decrease in noise, NS: the difference in noise between young and old individuals is not statistically significant. NP: the cell identity was not present in the dataset in sufficient amounts to perform the analysis. For most cell types, it can be concluded that there is little overlap between cell identity-specific noise measurements across datasets and methods

**Figure 4–Figure supplement 1.**
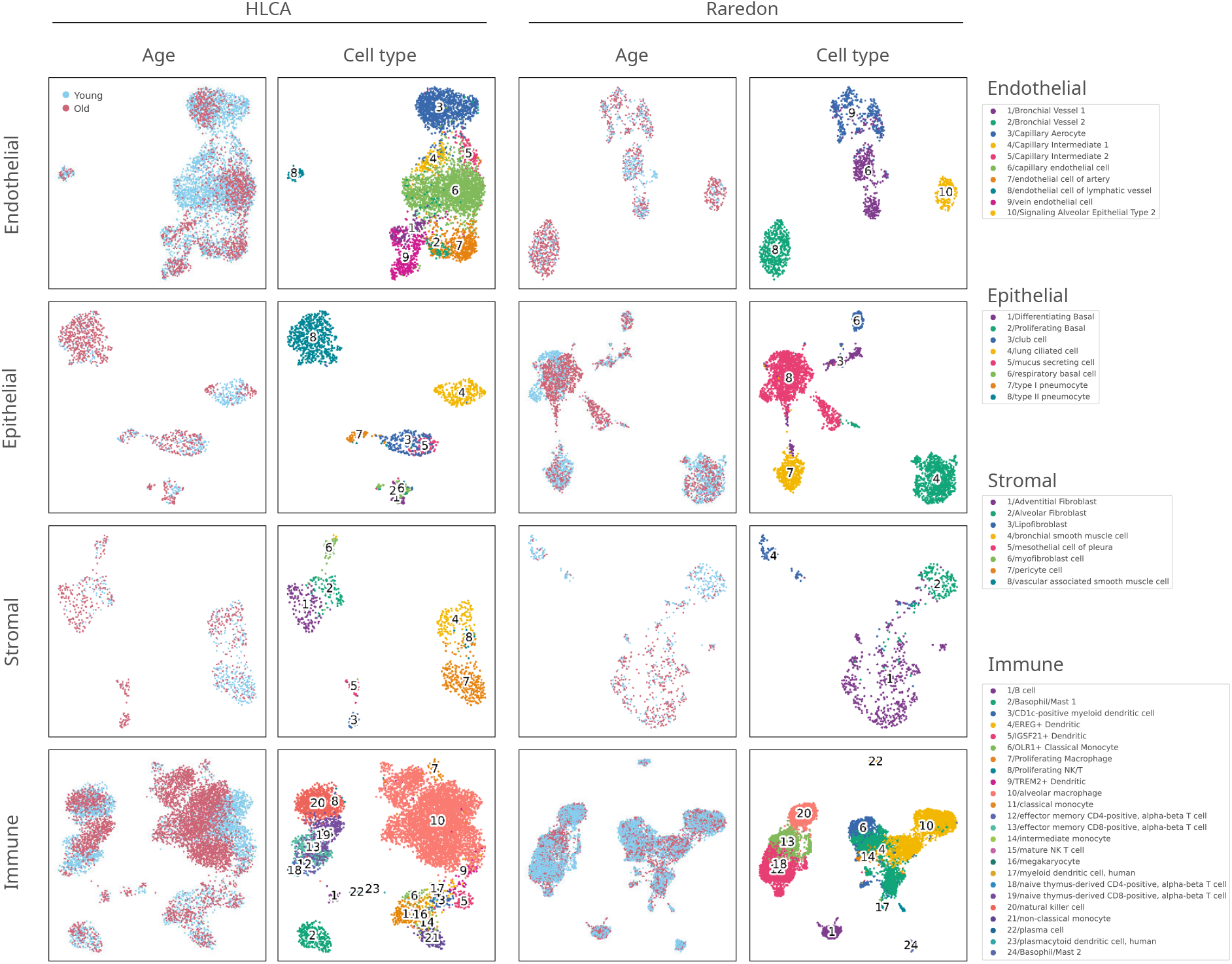
Composition of the two scRNAseq datasets of the human aging lung used in this figure. The UMAP plots with the age and cell type identity annotations are shown for each tissue compartment (endothelial, epithelial, stromal and immune) and each dataset separately.

**Figure 5–Figure supplement 1.**
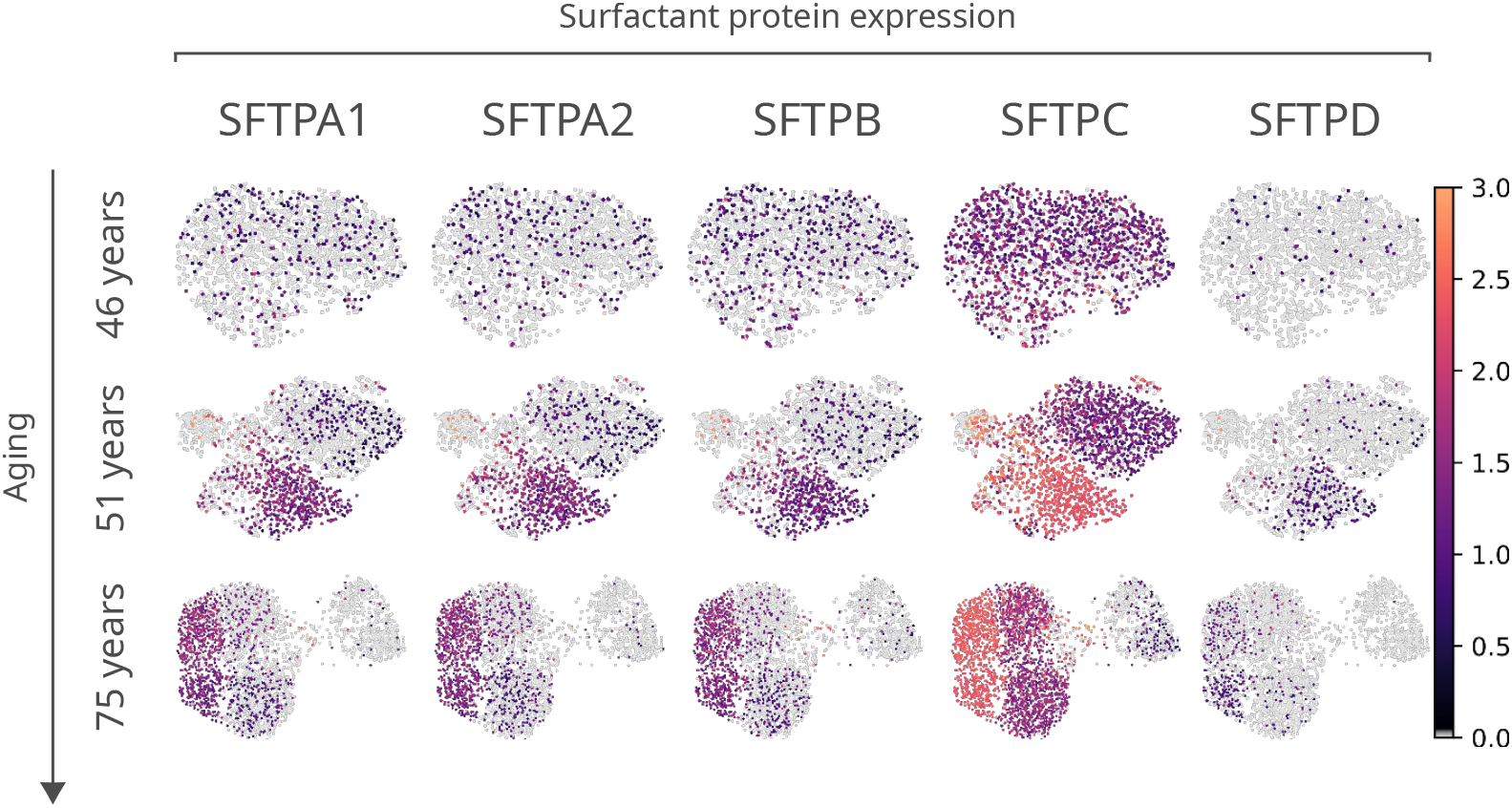
Expression of surfactant protein genes by human alveolar macrophages. Differential expression by alveolar macrophage cell clusters of the genes coding for surfactant proteins *SFTPA1, SFTPA2, SFTPB, SFTPC*, and *SFTPD* is shown for three donors (aged 46, 51 and 75) of the ***Travaglini et al***. (HLCA) dataset.

## Notes

### Competing Interest Statement

The authors have declared no competing interest.

## References

Almanzar N, Antony J, the Tabula Muris Consortium. A single-cell transcriptomic atlas characterizes ageing tissues in the mouse. Nature. 2020; doi:https://doi.org/10.1038/s41586-020-2496-1.

Angelidis I, Simon LM, Fernandez Ieea. An atlas of the aging lung mapped by single cell transcriptomics and deep tissue proteomics. Nature Communications. 2019; doi:https://doi.org/10.1038/s41467-019-08831-9.

Bacher R, Kendziorski C. Design and computational analysis of single-cell RNA-sequencing experiments. Genome Biology. 2016; 17:63. doi:https://10.1186/s13059-016-0927-y.

Bahar R, Hartmann CH, Rodriguez KA, Denny AD, Busuttil RA, Dollé MET, Calder RB, Chisholm GB, Pollock BH, Klein CA, et al. Increased cell-to-cell variation in gene expression in ageing mouse heart. Nature. 2006; 441(7096):1011–1014. doi:10.1038/nature04844.

Bezdek JC. Pattern Recognition with Fuzzy Objective Function Algorithms. USA: Kluwer Academic Publishers; 1981.

Blondel VD, Guillaume JL, Lambiotte R, Lefebvre E. Fast unfolding of communities in large networks. Journal of Statistical Mechanics: Theory and Experiment. 2008 oct; 2008(10):P10008. https://doi.org/10.1088/1742-5468/2008/10/p10008, doi:10.1088/1742-5468/2008/10/p10008.

Cagan A, Baez-Ortega A, Brzozowska N, Abascal F, Coorens THH, Sanders MA, Lawson ARJ, Harvey LMR, Bhosle S, Jones D, Alcantara RE, Butler TM, Hooks Y, Roberts K, Anderson E, Lunn S, Flach E, Spiro S, Januszczak I, Wrigglesworth E, et al. Somatic mutation rates scale with lifespan across mammals. Nature. 2022; 604:517–524. doi:10.1038/s41586-022-04618-z.

Changyou S, Wang L, Sen P. The eroding chromatin landscape of aging stem cells. Translational Medicine of Aging. 2020; 4:121–131. doi:10.1016/j.tma.2020.08.002.

Costa JPD, Vitorino R, Silva GM, Vogel C, Duarte AC, Rocha-Santos T. A synopsis on aging—Theories, mechanisms and future prospects. Ageing Research Reviews. 2016; 29:90–112. doi:10.1016/j.arr.2016.06.005.

Dunn JC. A Fuzzy Relative of the ISODATA Process and Its Use in Detecting Compact Well-Separated Clusters. Journal of Cybernetics. 1973; 3(3):32–57. https://doi.org/10.1080/01969727308546046, doi:10.1080/01969727308546046.

Duò A RM C S. A systematic performance evaluation of clustering methods for single-cell RNA-seq data. F1000Research. 2018; 7(1141). doi:https://doi.org/10.12688/f1000research.15666.2.

Enge M, Arda HE, Mignardi M, Beausang J, Bottino R, Kim SK, Quake SR. Single-Cell Analysis of Human Pancreas Reveals Transcriptional Signatures of Aging and Somatic Mutation Patterns. Cell. 2017; 171(2). doi:https://10.1016/j.cell.2017.09.004.

Fonseca Costa SS, Robinson-Rechavi M, Ripperger JA. Single-cell transcriptomics allows novel insights into aging and circadian processes. Brief Funct Genomics. 2021; 19(5-6):343–349. doi:10.1093/bfgp/elaa014.

Franceschi C, Bonafè M, Valensin S, Olivieri F, Luca MD, Ottaviani E, Benedictis GD. Inflamm-aging: An Evolutionary Perspective on Immunosenescence. Annals of the New York Academy of Sciences. 2000; 908(1):244–254. doi:10.1111/j.1749-6632.2000.tb06651.x.

Frasca D, Blomberg BB. Inflammaging decreases adaptive and innate immune responses in mice and humans. Biogerontology. 2015; 17(1):7–19. doi:10.1007/s10522-015-9578-8.

Freytag S, Tian L, Lönnstedt Iea. Comparison of clustering tools in R for medium-sized 10x Genomics single-cell RNA-sequencing data. F1000Research. 2018; 7. doi:https://doi.org/10.12688/f1000research.15809.1.

Gems D, de Magalhães JP. The hoverfly and the wasp: A critique of the hallmarks of aging as a paradigm. Ageing Res Rev. 2021; 70(101407). doi:10.1016/j.arr.2021.101407.

Gill D, Parry A, Santos F, Okkenhaug H, Todd CD, Hernando-Herraez I, Stubbs TM, Milagre I, Reik W. Multi-omic rejuvenation of human cells by maturation phase transient reprogramming. eLife. 2022; 11(e71624). doi:10.7554/eLife.71624.

Gladyshev VN. Aging: progressive decline in fitness due to the rising deleteriome adjusted by genetic, environmental, and stochastic processes. Aging Cell. 2016; 15(4). doi:10.1111/acel.12480.

Goldfarbmuren KC, Jackson ND, Sajuthi SP, Dyjack N, Li KS, Rios CL, Plender EG, Montgomery MT, Everman JL, Bratcher PE, et al. Dissecting the cellular specificity of smoking effects and reconstructing lineages in the human airway epithelium. Nature Communications. 2020; 11(1). doi:10.1038/s41467-020-16239-z.

Gupta K, Yadav P, Maryam S, Ahuja G, Sengupta D. Quantification of age-related decline in transcriptional homeostasis. J Mol Biol. 2021; 433(19):167179. doi:10.1016/j.jmb.2021.167179.

Ham L, Jackson M, Stumpf MP. Pathway dynamics can delineate the sources of transcriptional noise in gene expression. eLife. 2021; 10. doi:10.7554/elife.69324.

Hernando-Herraez I, Evano B, Stubbs T, Commere PH, Bonder MJ, Clark S, Andrews S, Tajbakhsh S, Reik W. Ageing affects DNA methylation drift and transcriptional cell-to-cell variability in mouse muscle stem cells. Nature Communications. 2019; 10(1). doi:10.1038/s41467-019-12293-4.

Izgi H, Han D, Isildak U, Huang S, Kocabiyik E, Khaitovich P, Somel M, Dönertas HM. Inter-tissue convergence of gene expression during ageing suggests age-related loss of tissue and cellular identity. eLife. 2022; 11. doi:10.7554/eLife.68048.

Joost S, Zeisel A, Jacob T, et al. Single-Cell Transcriptomics Reveals that Differentiation and Spatial Signatures Shape Epidermal and Hair Follicle Heterogeneity. Cell Systems. 2016; 3(3). doi:10.1016/j.cels.2016.08.010.

Kimmel JC, Penland L, Rubinstein ND, Hendrickson DG, Kelley DR, Rosenthal AZ. Murine single-cell RNA-seq reveals cell-identity- and tissue-specific trajectories of aging. Genome Research. 2019; 29(12):2088–2103. doi:10.1101/gr.253880.119.

Kirkwood TBL, Melov S. On the Programmed/Non-Programmed Nature of Ageing within the Life History. Current Biology. 2011; 21(18). doi:10.1016/j.cub.2011.07.020.

Kiselev VY, M ATSH. Challenges in unsupervised clustering of single-cell RNA-seq data. Nature Reviews Genetics. 2019; 20. doi:https://10.1038/s41576-018-0088-9.

Korsunsky I, Millard N, Fan J, Slowikowski K, Zhang F, Wei K, Baglaenko Y, Brenner M, Loh PR, Raychaudhuri S, et al. Fast, sensitive and accurate integration of single-cell data with Harmony. Nature Methods. 2019; 16(12):1289–1296. doi:10.1038/s41592-019-0619-0.

Kovacs EJ, Boe DM, Boule LA, Curtis BJ. Inflammaging and the lung. Clinics in Geriatric Medicine. 2017; 33(4):459–471. doi:10.1016/j.cger.2017.06.002.

Levy O, Amit G, Vaknin D, Snir T, Efroni S, Castaldi P, Liu YY, Cohen HY, Bashan A. Age-related loss of gene-to-gene transcriptional coordination among single cells. Nature Metabolism. 2020; 2(11):1305–1315. doi:10.1038/s42255-020-00304-4.

Li J, Zheng Y, Yan P, Song M, Wang S, Sun L, Liu Z, Ma S, Izpisua Belmonte JC, Chan P, Zhou Q, Zhang W, Liu GH, Tang F, Qu J. A single-cell transcriptomic atlas of primate pancreatic islet aging. Natl Sci Rev. 2020; 8(2):nwaa127. doi:10.1093/nsr/nwaa127.

Lu Y, Brommer B, Tian X, Krishnan A, Meer M, Wang C, Vera DL, Zeng Q, Yu D, Bonkowski MS, et al. Reprogramming to recover youthful epigenetic information and restore vision. Nature. 2020; 588(7836):124–129. doi:10.1038/s41586-020-2975-4.

Lu Y, Brommer B, Tian X, Krishnan A, Meer M, Wang C, Vera DL, Zeng Q, Yu D, Bonkowski MS, Yang JH, Zhou S, Hoffmann EM, Karg MM, Schultz MB, Kane AE, Davidsohn N, Korobkina E, Chwalek K, Rajman LA, et al. Reprogramming to recover youthful epigenetic information and restore vision. Nature. 2020; 588(7836):124–129. doi:10.1038/s41586-020-2975-4.

Lun A, McCarthy D, Marioni J. A step-by-step workflow for low-level analysis of single-cell RNA-seq data with Bioconductor. F1000Research. 2016; 5(2122). doi:10.12688/f1000research.9501.2.

López-Otín C, Blasco MA, Partridge L, Serrano M, Kroemer G. The Hallmarks of Aging. Cell. 2013; 153(6):1194–1217. doi:10.1016/j.cell.2013.05.039.

Ma S, Sun S, Geng L, Song M, Wang W, Ye Y, Ji Q, Zou Z, Wang S, He X, Li W, Rodriguez Esteban C, Long X, Guo G, Chan P, Zhou Q, Izpisua Belmonte JC, Zhang W, Qu J, Liu GH. Caloric restriction reprograms the single-cell transcriptional landscape of Rattus Norvegicus aging. Cell. 2020; 180:984–1001. doi:10.1016/j.cell.2020.02.008.

Ma S, Sun S, Li J, Fan Y, Qu J, Sun L, Wang S, Zhang Y, Yang S, Liu Z, et al. Single-cell transcriptomic atlas of primate cardiopulmonary aging. Cell Research. 2020; 31(4):415–432. doi:10.1038/s41422-020-00412-6.

Martinez-Jimenez CP, Eling N, Chen HC, Vallejos CA, Kolodziejczyk AA, Connor F, Stojic L, Rayner TF, Stubbington MJT, Teichmann SA, de la Roche M, Marioni JC, T OD. Aging increases cell-to-cell transcriptional variability upon immune stimulation. Science. 2019; 20(366). doi:https://10.1126/science.aah4115.

McCullagh P, Nelder JA. An outline of generalized linear models. Generalized Linear Models. 1989; p. 21–47. doi:10.1007/978-1-4899-3242-6_2.

McInnes L, Healy J, Saul N, Großberger L. UMAP: Uniform manifold approximation and projection. Journal of Open Source Software. 2018; 3(29):861. doi:10.21105/joss.00861.

McQuattie-Pimentel AC, Ren Z, Joshi N, Watanabe S, Stoeger T, Chi M, Lu Z, Sichizya L, Aillon RP, Chen CI, et al. The lung microenvironment shapes a dysfunctional response of alveolar macrophages in aging. Journal of Clinical Investigation. 2021; 131(4). doi:10.1172/jci140299.

Mendenhall AR, Martin GM, Kaeberlein M, Anderson RM. Cell-to-cell variation in gene expression and the aging process. Geroscience. 2021; 43(1):181–196. doi:10.1007/s11357-021-00339-9.

Misharin AV, Morales-Nebreda L, Reyfman PA, Cuda CM, Walter JM, McQuattie-Pimentel AC, Chen CI, Anekalla KR, Joshi N, Williams KJN, et al. Monocyte-derived alveolar macrophages drive lung fibrosis and persist in the lung over the life span. Journal of Experimental Medicine. 2017; 214(8):2387–2404. doi:10.1084/jem.20162152.

Mishra S, Srivastava D, Kumar V. Improving gene network inference with graph wavelets and making insights about ageing-associated regulatory changes in lungs. Brief Bioinform. 2021; 22(4):bbaa360. doi:10.1093/bib/bbaa360.

Moignard V, Woodhouse S, Haghverdi L, et al. Decoding the regulatory network of early blood development from single-cell gene expression measurements. Nature Biotechnology. 2015; 33(3). doi:10.1038/nbt.3154.

Munkres J. Algorithms for the Assignment and Transportation Problems. Journal of the Society of Industrial and Applied Mathematics. 1957; 5(1).

Nalapareddy K, Zheng Y, Geiger H. Aging of intestinal stem cells. Stem Cell Reports. 2022; 17(4):734–740. doi:10.1016/j.stemcr.2022.02.003.

Nikopoulou C, Parekh S, Tessarz P. Ageing and sources of transcriptional heterogeneity. Biological Chemistry. 2019; 400(7):867–878. doi:10.1515/hsz-2018-0449.

Oh J, Lee YD, Wagers AJ. Stem cell aging: Mechanisms, regulators and therapeutic opportunities. Nature Medicine. 2014; 20(8):870–880. doi:10.1038/nm.3651.

Olah M, Patrick E, Villani AC, Xu J, White CC, Ryan KJ, Piehowski P, Kapasi A, Nejad P, Cimpean M, et al. A transcriptomic atlas of aged human microglia. Nature Communications. 2018; 9(1). doi:10.1038/s41467-018-02926-5.

Oliviero G, Kovalchuk S, Rogowska-Wrzesinska A, Schwämmle V, Jensen ON. Distinct and diverse chromatin proteomes of ageing mouse organs reveal protein signatures that correlate with physiological functions. eLife. 2022; 11(e73524). doi:10.7554/eLife.73524.

Park JE, K P K M, Teichmann SA. Fast Batch Alignment of Single Cell Transcriptomes Unifies Multiple Mouse Cell Atlases into an Integrated Landscape. bioRxiv. 2018; https://www.biorxiv.org/content/early/2018/08/22/397042, doi:10.1101/397042.

Paul F, Arkin Y, Giladi A, et al. Transcriptional Heterogeneity and Lineage Commitment in Myeloid Progenitors. Nature Biotechnology. 2015; 163(7). doi:10.1016/j.cell.2015.11.013.

Pawelec G. Hallmarks of human “immunosenescence”: adaptation or dysregulation? Immunity & ageing. 2012; 9(1). doi: doi.org/10.1186/1742-4933-9-15.

Pisco A. Tabula muris senis. Processed files (to use with scanpy).. 2020; doi:10.6084/m9.figshare.12654728.v1.

Pálovics R, Keller A, Schaum N, Tan W, Fehlmann T, Borja M, Kern F, Bonanno L, Calcuttawala K, Webber J, McGeever A, Consortium TTM, Luo J, Pisco AO, Karkanias J, Neff NF, Darmanis S, Quake SR, Wyss-Coray T. Molecular hallmarks of heterochronic parabiosis at single-cell resolution. Nature. 2022; 603:309–314. doi:10.1038/s41586-022-04461-2.

Raredon MSB, Adams TS, Suhail Y, Schupp JC, Poli S, Neumark N, Leiby KL, Greaney AM, Yuan Y, Horien C, et al. Single-cell connectomic analysis of adult mammalian lungs. Science Advances. 2019; 5(12). doi:10.1126/sciadv.aaw3851.

Raser JM, O’Shea EK. Noise in gene expression: origins, consequences, and control. Science. 2005; 309(5743):2010–2013. doi:10.1126/science.1105891.

Rhoades NS, Davies M, Lewis SA, Cinco IR, Kohama SG, Bermudez LE, Winthrop KL, Fuss C, Mattison JA, Spindel ER, Messaoudi I. Functional, transcriptional, and microbial shifts associated with healthy pulmonary aging in rhesus macaques. Cell Reports. 2022; 39(110725). doi:10.1016/j.celrep.2022.110725.

Salminen A. Feed-forward regulation between cellular senescence and immunosuppression promotes the aging process and age-related diseases. Ageing Res Rev. 2021; 67(101280). doi:10.1016/j.arr.2021.101280.

Salzer MC, Lafzi A, Berenguer-Llergo A, Youssif C, Castellanos A, Solanas G, Peixoto FO, Attolini CSO, Prats N, Aguilera M, et al. Identity Noise and Adipogenic Traits Characterize Dermal Fibroblast Aging. Cell. 2018; 175(6). doi:10.1016/j.cell.2018.10.012.

Satija R, Farrell D J A Gennert, Schier AF, Regev A. Spatial reconstruction of single-cell gene expression data. Nature Biotechnology. 2015; doi:https://doi.org/10.1038/nbt.3192.

Schaum N, Lehallier B, Hahn O, Pálovics R, Hosseinzadeh S, Lee SE, Sit R, Lee DP, Morán Losada P, Zardeneta ME, Fehlmann T, Webber JT, McGeever A, Calcuttawala K, Zhang H, Berdnik D, Mathur V, Tan W, Zee A, Tan M, et al. Ageing hallmarks exhibit organ-specific temporal signatures. Nature. 2020; 583:309–314. doi:10.1038/s41586-022-04461-2.

Schiller HB, Montoro DT, Simon LM, Rawlins EL, Meyer KB, Strunz M, Vieira Braga FA, Timens W, Koppelman GH, Budinger GR, et al. The Human Lung Cell Atlas: A high-resolution reference map of the human lung in health and disease. American Journal of Respiratory Cell and Molecular Biology. 2019; 61(1):31–41. doi:10.1165/rcmb.2018-0416tr.

Schmeer C, Kretz A, Wengerodt D, Stojiljkovic MA, Witte OW. Dissecting Aging and Senescence-Current Concepts and Open Lessons. Cells. 2019; 8(11). doi:10.3390/cells8111446.

Schneider JL, Rowe JH, Garcia-de Alba C, Kim CF, Sharpe AH, Haigis MC. The aging lung: Physiology, disease, and immunity. Cell. 2021; 184(8):1990–2019. doi:10.1016/j.cell.2021.03.005.

Searle SR, Speed FM, Milliken GA. Population Marginal Means in the Linear Model: An Alternative to Least Squares Means. The American Statistician. 1980; 34(4):216–221. doi:10.1080/00031305.1980.10483031.

Seninge L. scoreCT: Automated Cell Type Annotation. GitHub repository. 2020;.

Sikkema L, Strobl D, Zappia L, Madissoon E, Markov NS, Zaragosi L, Ansari M, Arguel M, Apperloo L, Bécavin C, Berg M, Chichelnitskiy E, Chung M, Collin A, Gay ACA, Hooshiar Kashani B, Jain M, Kapellos T, Kole TM, Mayr C, et al. An integrated cell atlas of the human lung in health and disease. bioRxiv. 2022; doi:10.1101/2022.03.10.483747.

Solé-Boldo L, Raddatz G, Schütz Sea. Single-cell transcriptomes of the human skin reveal age-related loss of fibroblast priming. Commun Biol. 2020; 3(188). doi:https://doi.org/10.1038/s42003-020-0922-4.

Takemon Y, Chick JM, Gerdes Gyuricza I, Skelly DA, Devuyst O, Gygi SP, Churchill GA, Korstanje R. Proteomic and transcriptomic profiling reveal different aspects of aging in the kidney. eLife. 2021; 10. doi:10.7554/elife.62585.

Tasic B, Menon V, Nguyen TN, Kim TK, Jarsky T, Yao Z, Levi B, Gray LT, Sorensen SA, Dolbeare T, et al. Adult mouse cortical cell taxonomy revealed by single cell transcriptomics. Nature Neuroscience. 2016; 19(2):335–346. doi:10.1038/nn.4216.

Traag VA, Waltman L, van Eck NJ. From Louvain to Leiden: guaranteeing well-connected communities. Scientific Reports. 2019; 9. doi:https://doi.org/10.1038/s41598-019-41695-z.

Trapnell C. Defining cell types and states with single-cell genomics. Genome Research. 2015; 25(10):1491–1498. doi:10.1101/gr.190595.115.

Travaglini KJ, Nabhan AN, Penland L, Sinha R, Gillich A, Sit RV, Chang S, Conley SD, Mori Y, Seita J, et al. A molecular cell atlas of the human lung from single-cell RNA sequencing. Nature. 2020; 587(7835):619–625. doi:10.1038/s41586-020-2922-4.

Uyar B, Palmer D, Kowald A, Murua-Escobar H, Barrantes I, Möller S, Akalin A, Fuellen G. Singlecell analyses of aging, inflammation and senescence. Ageing Res Rev. 2020; 64(101156). doi:https://10.1016/j.arr.2020.101156.

Vijg J. From DNA damage to mutations: All roads lead to aging. Ageing Res Rev. 2021; 68(101316). doi:10.1016/j.arr.2021.101316.

Waltman L, van Eck NJ, Traag VA. A smart local moving algorithm for large-scale modularity-based community detection. European Physical Journal B. 2013; 86.

Warren LA, Rossi DJ, Schiebinger GR, Weissman IL, Kim SK, Quake SR. Transcriptional instability is not a universal attribute of aging. Aging Cell. 2007; 6(6):775–782. doi:10.1111/j.1474-9726.2007.00337.x.

Williamson EJ, Walker AJ, Bhaskaran K, Bacon S, Bates C, Morton CE, Curtis HJ, Mehrkar A, Evans D, Inglesby P, et al. Factors associated with COVID-19-related death using OpenSAFELY. Nature. 2020; 584(7821):430–436. doi:10.1038/s41586-020-2521-4.

Wolf FA, p A, Theis FJ. SCANPY: large-scale single-cell gene expression data analysis. Genome Biology. 2018; doi: https://doi.org/10.1186/s13059-017-1382-0.

Ximerakis M, Lipnick SL, Innes BT, Simmons SK, Adiconis X, Dionne D, Mayweather BA, Nguyen L, Niziolek Z, Ozek C, Butty VL, Isserlin R, Buchanan SM, Levine SS, Regev A, Bader GD, Levin JZ, Rubin LL. Single-cell transcriptomic profiling of the aging mouse brain. Nat Neurosci. 2019; 22(10). doi:https://10.1038/s41593-019-0491-3.

Yang JH, Griffin PT, Vera DL, Apostolides JK, Hayano M, Meer MV, Salfati EL, Su Q, Munding EM, Blanchette M, Bhakta M, Dou Z, Xu C, Pippin JW, Creswell ML, O’Connell BL, Green RE, Garcia BA, Berger SL, Oberdoerffer P, et al. Erosion of the Epigenetic Landscape and Loss of Cellular Identity as a Cause of Aging in Mammals. bioRxiv. 2019; https://www.biorxiv.org/content/early/2019/10/19/808642, doi:10.1101/808642.

Yousefzadeh MJ, Flores RR, Zhu Y, Schmiechen ZC, Brooks RW, Trussoni CE, Cui Y, Angelini L, Lee KA, McGowan SJ, Burrack AL, Wang D, Dong Q, Lu A, Sano T, O’Kelly R, McGuckian CA, Kato JI, Bank MP, Wade EA, et al. An aged immune system drives senescence and ageing of solid organs. Nature. 2021; 594(7861):100–105. doi:10.1038/s41586-021-03547-7.

Zhang W, Zhang S, Yan P, Ren J, Song M, Li J, Lei J, Pan H, Wang S, Ma X, et al. A single-cell transcriptomic landscape of primate arterial aging. Nature Communications. 2020; 11(1). doi:10.1038/s41467-020-15997-0.

Zhu L, Lei J, Klei L, Devlin B, Roeder K. Semisoft clustering of single-cell data. Proceedings of the National Academy of Sciences. 2019; 116(2):466–471. https://www.pnas.org/content/116/2/466, doi:10.1073/pnas.1817715116.

Zou Z, Long X, Zhao Q, Zheng Y, Song M, Ma S, Jing Y, Wang S, He Y, Esteban CR, et al. A Single-Cell Transcriptomic Atlas of Human Skin Aging. Developmental Cell. 2021; 56(3). doi:10.1016/j.devcel.2020.11.002.

